# Predicting direct physical interactions in multimeric proteins with deep learning

**DOI:** 10.1101/2021.11.09.467949

**Authors:** Mu Gao, Davi Nakajima An, Jerry M. Parks, Jeffrey Skolnick

## Abstract

Accurate descriptions of protein-protein interactions are essential for understanding biological systems. Remarkably accurate atomic structures have been recently computed for individual proteins by AlphaFold2 (AF2). Here, we demonstrate that the same neural network models from AF2 developed for single protein sequences can be adapted to predict the structures of multimeric protein complexes without retraining. In contrast to common approaches, our method, AF2Complex, does not require paired multiple sequence alignments. It achieves higher accuracy than some complex protein-protein docking strategies and provides a significant improvement over AF-Multimer, a new development of AlphaFold for multimeric proteins. Moreover, we introduce metrics for predicting direct protein-protein interactions between arbitrary protein pairs and validate AF2Complex on some challenging benchmark sets and the *E. coli* proteome. Lastly, using the cytochrome *c* biogenesis system I as an example, we present high-confidence models of three sought-after assemblies formed by eight members of this system.

## Introduction

AlphaFold2 (AF2), a deep learning approach developed by DeepMind for predicting protein structure given a sequence, has greatly advanced protein structure prediction[1, 2]. In CASP14, a blind test for protein structure prediction, AF2 achieved remarkable performance when assessed on the individual domains of target protein sequences[1]. Because deep learning is a data-driven approach, two key factors contributing to the success of AF2 are the completeness of the structural space of single-domain proteins and an abundance of sequences in sequence databases[3]. Together, these factors have made it possible to train sophisticated neural network models for accurate protein structure prediction.

In addition to single-domain targets, AF2 also delivered excellent results on multi-domain proteins[1] and has been applied to such proteins in several model organisms[2]. Considering that intra-protein domain-domain interactions are not physically different from inter-protein interactions, these AF2 results are very intriguing. They hint that, in principle, AF2 could be repurposed to predict whether a pair of proteins interact and if so, to predict the quaternary structure of the resulting protein complex. After all, many proteins that form complexes in prokaryotes are fused into long, single-chain, multi-domain proteins in eukaryotes[4]. The same physical forces that drive protein folding are also responsible for protein-protein associations[5]. Moreover, it is known that the structural space of protein-protein interfaces, the regions where direct physical contacts are found between protein partners forming a complex, is quite small[6]. Taken together, it is very likely that the neural network models trained for single-chain structure prediction have already learned the representations necessary to model protein complexes made of multiple single-chain proteins[3]. Indeed, some examples of using such a neural network model to predict complex structures were demonstrated with RoseTTAFold[7], whose design was inspired by AF2, even though its examples were likely present in its own model training set.

Until now, conventional approaches for predicting the structure of protein complexes include a docking component[8–11], which is limited by force-field accuracy and the completeness of conformational space sampling. A second class of approaches is template-based methods, which utilize experimentally resolved multimeric structures[12–16]. More recent methods incorporate inter-protein residue-residue contact predictions obtained via co-evolutionary analysis[17] and a deep learning algorithm[18].

Can AF2 be adapted to predict the structure of a protein complex? After the release of AF2, efforts immediately began to seek an answer. The first such study involved simply connecting two protein sequences with a poly-glycine linker, thus converting it into a single-chain structure prediction problem[19]. A much better solution is to modify the *“residue_index”* feature used by AF2, which eliminates the need for a covalent linker that likely creates artifacts[20, 21]. Meanwhile, studies have been carried out whereby models of single proteins generated with AF2 are used with docking methods. They are based on the idea that AF2 generates high-quality monomeric models that could improve the chance of native-like poses in docking[22]. One issue with these studies, as some authors pointed out, is that the benchmark set tested includes protein structures used to train the AF2 deep learning models. Although the AF2 models were not trained on protein complex structures, the use of the holo monomers in training compromises rigor because AF2 likely provides an “observed” holo-structure for docking.

Going beyond the prediction of the structure of the protein complex given that the proteins interact, are more fundamental but more challenging questions: Can AF2 be adapted to predict protein-protein interactions given an arbitrary pair of protein sequences, and more generally, can it identify higher order protein complexes? Several high-throughput experimental techniques have been designed to identify interacting protein partners[23–26], but their results are far from complete and are often mutually inconsistent. Computationally, template-based approaches have been used[27], but they are limited to the detection of homologs. Combining standard protein-protein docking methods with co-evolutionary signals[28] or even with deep learning models[29] has also been carried out on full proteomes. These are powerful approaches, but they rely on paired multiple sequence alignments (MSAs) as inputs. Generating paired MSAs requires the identification of orthologous sequences across species, which is impractical in many cases because it is confounded by the presence of paralogs in eukaryotes, protein cross-talk in disease pathways, and novel pathogen-host interactions. After all, one main consequence of evolution is the diversification of protein functions by producing paralogs[30]. These paralogs may interact with different partners without using a conserved interaction mode. Therefore, it is highly desirable to develop an approach that is not dependent on paired multiple sequence alignments.

In this study, using multiple test sets and *without* using paired sequence alignments, we demonstrate that AF2 can be adapted to predict both the presence of protein-protein interactions and the corresponding quaternary structures. Although our tests are primarily conducted on dimers, the method, AF2Complex, can be applied to higher-order oligomers, and we show examples of such. Critically, it is necessary to devise proper metrics to estimate the confidence of a predicted protein complex model. By adapting metrics previously introduced for comparing the similarity of protein-protein interfaces[31], we introduce new metrics for assessing the likelihood of protein-protein interactions. When AF2Complex was applied to a previously defined “gold standard” interaction set in *E. coli* [32], it found that many protein pairs are likely due to associations in large assemblies that are not necessarily in direct physical contact. Finally, we apply AF2Complex to make novel predictions on sought-after assemblies of a cytochrome *c* biogenesis system[33, 34].

## Results

An overview of AF2Complex is illustrated in Fig. 1 with the details in Methods. Given query sequences of a target protein complex, the input features for each query are first collected by applying the original AF2 data pipeline. Then, AF2Complex assembles the individual monomer features for complex structure prediction. Among the input features, the most critical are the MSAs, which are obtained by extending each monomeric alignment sequence to the full complex length with gap paddings. Correspondingly, to mark separate peptide chains we sequentially increase the residue index feature of the second or later monomer(s) by a large number. The structural templates of monomer sequences are also re-indexed accordingly. If the input contains multiple copies of the same sequence, i.e., a homo-oligomer, it is treated as if they were heterogeneous sequences. In this way, one can readily reuse pre-computed features for individual sequences, e.g., from proteomes of species, for protein-protein interaction screening without any extra step such as MSA pairing. The input features for the putative complex are then separately supplied to AF2 DL models, and the resulting structure models retained for analysis. Finally, the likelihood of complex formation is assessed by two metrics: the interface-score and the predicted interface TM-score (piTM), both of which evaluate the confidence of the predicted protein-protein interface if found in an assessed model (see Methods). Each of these two scores ranges from 0 to 1, where a higher score indicates higher confidence.

**Fig. 1.**
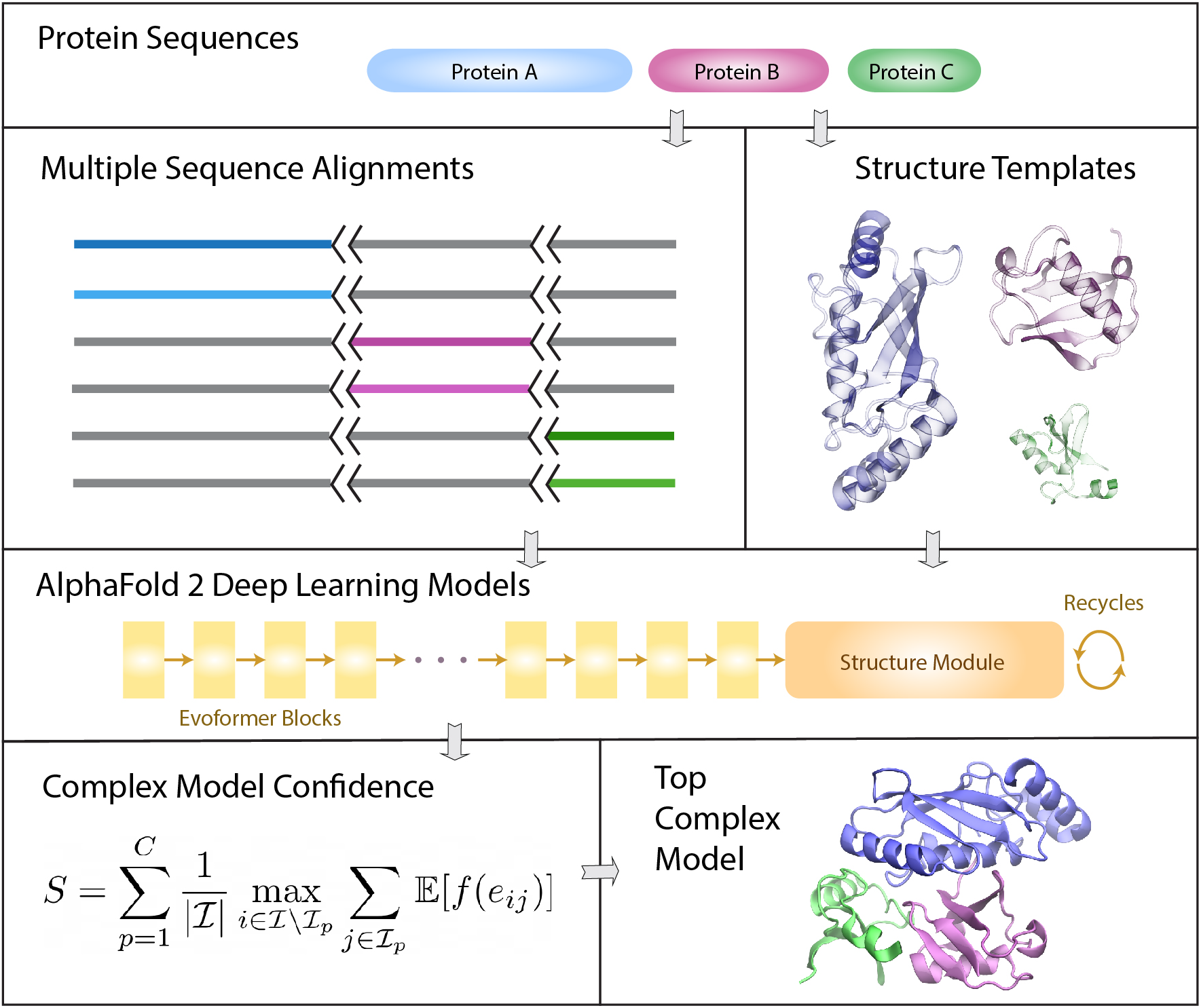
Overview of the AF2Complex workflow. The multiple sequence alignments of query protein sequences A (blue), B (purple), and C (green) are joined together by padding gaps (grey) in the MSA regions belonging to other proteins, and the short black lines represent an increase in the residue index to distinguish separate protein chains. Structure templates for individual proteins are also retrieved from the Protein Data Bank. Using these sequence and template features, an AF2 DL model generates a complex model after multiple recycles. The interface residues between proteins in the final complex model are then identified and their interface-score *S* is calculated to rank model confidence (see Methods).

We note that AF2Complex was originally based on the five monomer DL models of AF2. After the release of AF-Multimer[35], additionally AF2Complex was adapted to take five multimer DL models for predictions using unpaired MSAs. Throughout this work, without note, we present structural models obtained with AF2’s monomer DL models, rather than with AF-Multimer models.

### Accurate predictions on some CASP14 multimeric targets

We first applied AF2Complex to the multimeric targets of CASP14[36, 37]. To simulate a CASP14 prediction scenario (see Methods), the input feature predictions described below were obtained by using databases available prior to May 15, 2020, CASP14’s starting date. Because the structures of many assembly targets have not been released to the public, it is impossible to conduct a reliable statistical analysis of our predictions. However, they do showcase the potential of AF2Complex.

Fig. 2 displays the results of AF2Complex on some challenging targets. The first example, H1072, is a heterotetramer (stoichiometry A_2_B_2_) consisting of two copies of two coiled-coil protein sequences[39]. Despite the simple topologies and the availability of an experimental structure for one monomer, H1072 is a difficult target. No group participating in the CASP14 competition provided a correct model of the complex[36]. In contrast, the top model by AF2Complex achieved a remarkable TM-score[40] of 0.90 when superimposed onto the experimental structure (Fig. 2a). The second example, H1065, is a challenging heterodimer as one component lacks a homolog in the PDB[36]. In this case, AF2Complex generated a highly accurate complex model with a TM-score of 0.94 (Fig. 2b). The interface similarity score (IS-score), which was designed to evaluate dimeric protein-protein interfaces[41], is 0.60 with a significant *P*-value of 2×10^-20^. The third example, T1070o, is a homo-trimer with intertwined β-sheets at the N-termini (Fig. 2c). Although we could not evaluate the overall complex structure because its experimental structure is unavailable, we were able to evaluate a monomeric structure that contains a free-modeling domain target (T1070-D1). If we extract this monomeric domain from our top complex model and compare it to the native structure, our model for T1070-D1 yields a TM-score of 0.74, which is a significant improvement over 0.62 by AF2 in its official CASP14 assessment. This example indicates that, by modeling the entire homo-oligomeric target complex, one may obtain a structural model with higher quality, especially for an intertwined oligomer.

**Fig. 2.**
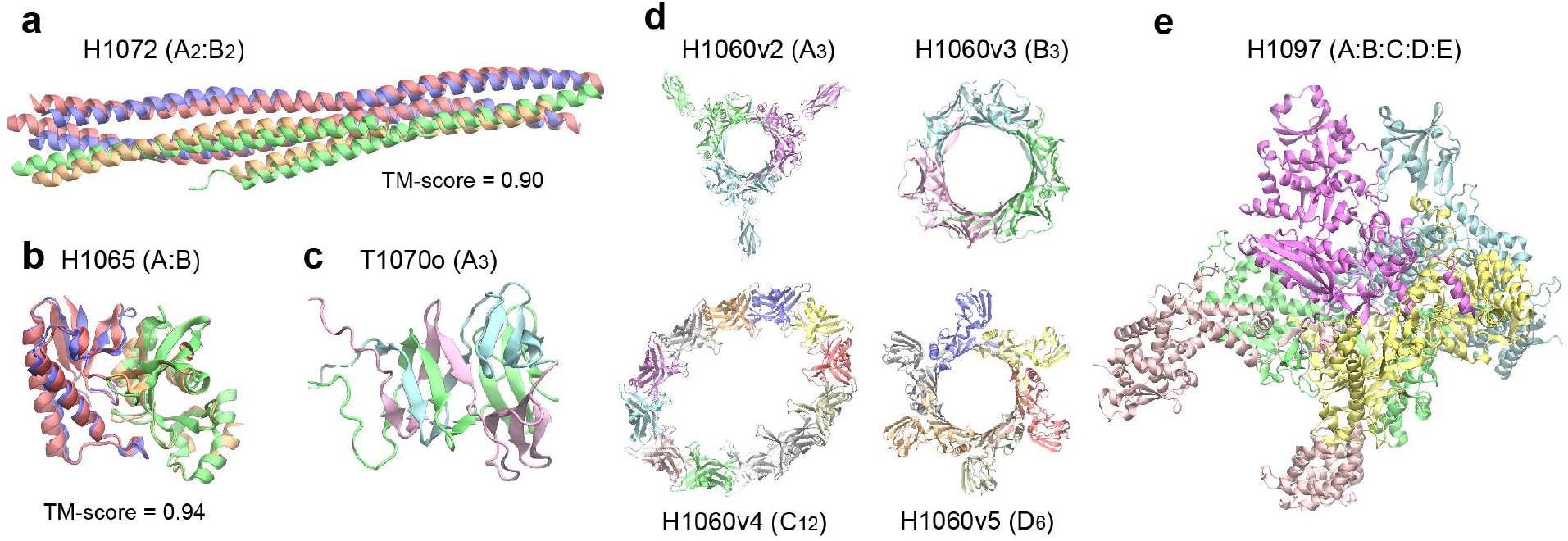
Top complex models generated by AF2Complex for selected CASP14 assembly targets. Each target is labeled with its target name, e.g., H1072, followed by its stoichiometry in parentheses, e.g., A_2_:B_2_. For targets with available experimental structure coordinates, the TM-score between the model and experimental structure is provided. For other structures only an image of the predicted model is given. Models are colored red and green, and experimental structures are in blue and gold. (**a**) SYCE2-TEX12 delta-Ctip complex. (**b**) N4-cytosine methyltransferase. (**c**) G3M192 from *Escherichia* virus CBA120. Only the N-terminal domains, which have an intertwined complex structure, are shown from a model of the full trimer. (**d**) Four rings from the T5 phage tail subcomplex. (**e**) DNA-directed RNA polymerase from *Bacillus* phage AR9. All images were generated with VMD [38].

The last two targets are from large molecular machines whose full experimental coordinates are unavailable. One of them, H1060, is part of a T5 phage tail assembly[36, 37]. The total size of this complex target is 6,582 residues, which is too large to be modeled in full. However, it is possible to model its four ring-like sub-structures, which are formed by 3 to 12 copies of four distinct monomers. AF2Complex returns models of these four rings at highly confident interface-scores ranging from 0.56 to 0.83 (Fig. 2d). The most challenging one is the 12-membered ring, for which our model forms an ellipse instead of the expected circular structure. Using AF-Multimer DL models, we obtained a single model with the expected C12 symmetry using unpaired MSAs but failed to build a physical model with paired MSAs (see SI). The last example, H1097, is a DNA-directed RNA polymerase from *Bacillus* phage AR9. It is composed of five hetero-monomers, totaling 2,682 amino acids. AF2Complex generates a highly confident model with an interface-score of 0.79 (Fig. 2e). Given that there are quite a few RNA polymerase structures in the current PDB, perhaps this result is not surprising. But the fact that AF2Complex can produce a model without paired MSAs in this case strongly indicates that paired MSAs might not be essential.

### Significantly higher accuracy over docking-centric approaches and improvement on AF-Multimer

Next, we conducted a benchmark test using 17 heterodimers released after Apr 30, 2018, which was the cutoff date of PDB structures collected for training the AF2 models. This set, named CP17, was curated for assessing various docking-centric strategies in a recent study[22]. One such strategy is to build a complex model using the ColabFold version of AF2[20], then split the monomers from the predicted AF complex and use the ClusPro[42] docking method to generate complex models. This strategy yields an acceptable or better top-ranked model for fewer than half of the targets. By comparison, the overall top models from AF2Complex are acceptable or better (see Methods) in 15 of 17 (88%) cases, and 13 (76%) models are of medium or high quality according to the DockQ score[43] (Fig. 3a, Supplementary Table 1), with a mean of 0.62, a dramatic improvement from 0.25 of the docking-centric approach. Another complex, docking-centric strategy increases the mean of DockQ score from 0.25 to 0.47[22]. Nevertheless, the improvement still falls behind AF2Complex on the same set (Fig. 3b). However, in one of the two cases when AF2Complex failed, the combined strategy resulted in a high-quality model. In these two failed cases, each has one monomer with only single-digit depth in its monomeric MSAs, which may explain these failures.

**Fig. 3.**
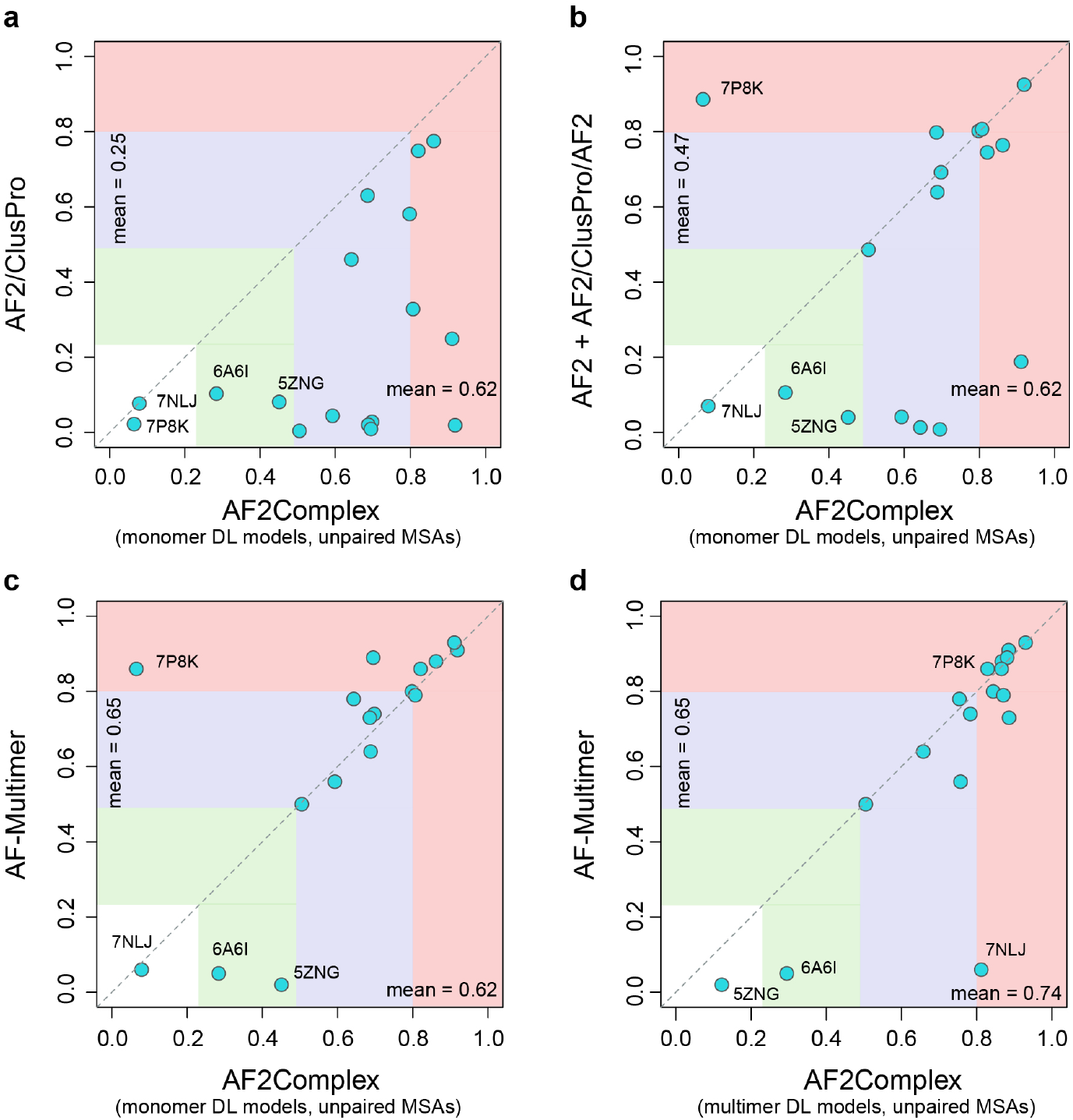
Comparison of AF2Complex and three alternative approaches on the CP17 data set. The coordinates of the circles correspond to the DockQ scores of the top overall models from each approach versus AF2Complex. (**a**) AF2 models docked by ClusPro, (**b**) Docking models refined by AF2, plus additional complex models obtained by running AF2 on paired MSAs according to ref [22], and (**c** & **d**) AlphaFold-Multimer [35]. The AF2 deep learning models trained for the prediction of monomeric protein structures, denoted as “monomer DL models”, were employed by AF2Complex in (A-C), and the AF-Multimer deep learning models, denoted as “multimer DL models”, were applied with AF2Complex in (d). All MSA inputs to AF2Complex are unpaired as described in Methods. Vertical and horizontal blocks represent the regions of incorrect (white), acceptable (green), medium (blue), and high-quality (red) complex models according to the DockQ score (see Methods).

Moreover, using AF2 monomer DL models, AF2Complex performs quite close to AF-Multimer[35] on the CP17 set (Fig. 3c), with a mean DockQ score of 0.62 versus 0.65. The mean score of AF2Complex increases to 0.74 using AF-Multimer DL models (Fig. 3d), thanks to unpaired MSAs, increased recycles and the interface-score metric (Supplementary Table 2). By combining the overall best model from AF2Complex runs using either monomer and multimer DL models of AF, we obtain acceptable or better models for all 17 targets.

To address the small target size of CP17, a large-scale benchmark study was further carried out on 1,192 dimers and 593 oligomers (see Methods and SI). On 440 heterodimers obtained using the same DL models, AF2Complex performs significantly better than AF-Multimer, albeit at smaller advantage (median/mean 0.69/0.56 versus 0.65/0.55, *p*-value = 5×10^-3^, see SI and Fig. S2), whereas their performance on the homodimer set is comparable (Fig. S2). AF2Complex further improves (median/mean 0.70/0.57 on the heterodimers, *p*-value = 6×10^-4^) if the overall top ranked model of both monomer and multimer DL runs are assessed. It must be pointed that AF-Multimer (version 2.1.1) may yield unphysical models with severe clashes for a complex, especially for large oligomeric targets including homo-oligomers. In contrast, AF2Complex mitigates this issue with unpaired MSAs using the same multimer DL models and deals much more effectively with unpaired MSAs and the original monomer DL models of AF2 (SI and Fig. S3).

### Predicting physical interactions among arbitrary protein pairs

Having been rather successful, an obviously tantalizing question is: Can this approach be applied to predict direct protein-protein interactions for an arbitrary pair of proteins? To answer this question, we devised a new test using the 34 unique protein sequences from the CP17 set. The goal was to find the 17 true interacting pairs given in CP17 from the 561 all-against-all pairwise combinations. Here, we naively assumed that all protein pairs other than the CP17 pairs are non-interacting, and any hit above a cutoff value of a metric adopted for evaluation is a false positive. Fig. 4 shows the results by using four different metrics to predict protein-protein interactions. Note that the model predictions for all pairs were carried out under exactly the same configuration in AF2Complex runs. Overall, both the interface-score and the piTM-score demonstrate a clear advantage over the other two metrics, the pTM-score and pLDDT-score of AF2[1]. Because we expect that most pairs of proteins are non-interacting, we focus on the regime of low false positive rate (i.e., FP < 0.1) in the receiver operating characteristic (ROC) curve. The normalized area under the curve (AUC) of this plot, AUC0.1, is 0.72 and 0.69 for interface-score and piTM, versus 0.49 and 0.10 for pTM and pLDDT, respectively. For reference, random guessing yields an AUC0.1 of 0.05. As expected, pLDDT is not ideal for evaluating protein complex models because it was designed for single domain evaluation. Although the pTM metric is much more discriminating than pLDDT, it is still much worse than interface-score or piTM in this regard. The same trend is also displayed in the precision-recall plot, whereby we achieved ~45% recall (equivalent to the true positive rate) at ~90% precision, and the recall increases to ~70% at ~45% precision. Correspondingly, the interface-score and piTM values are 0.55/0.59 and 0.45/0.50, respectively. Overall, the results encouragingly conclude that AF2Complex can be used to predict protein-protein interactions.

**Fig. 4.**
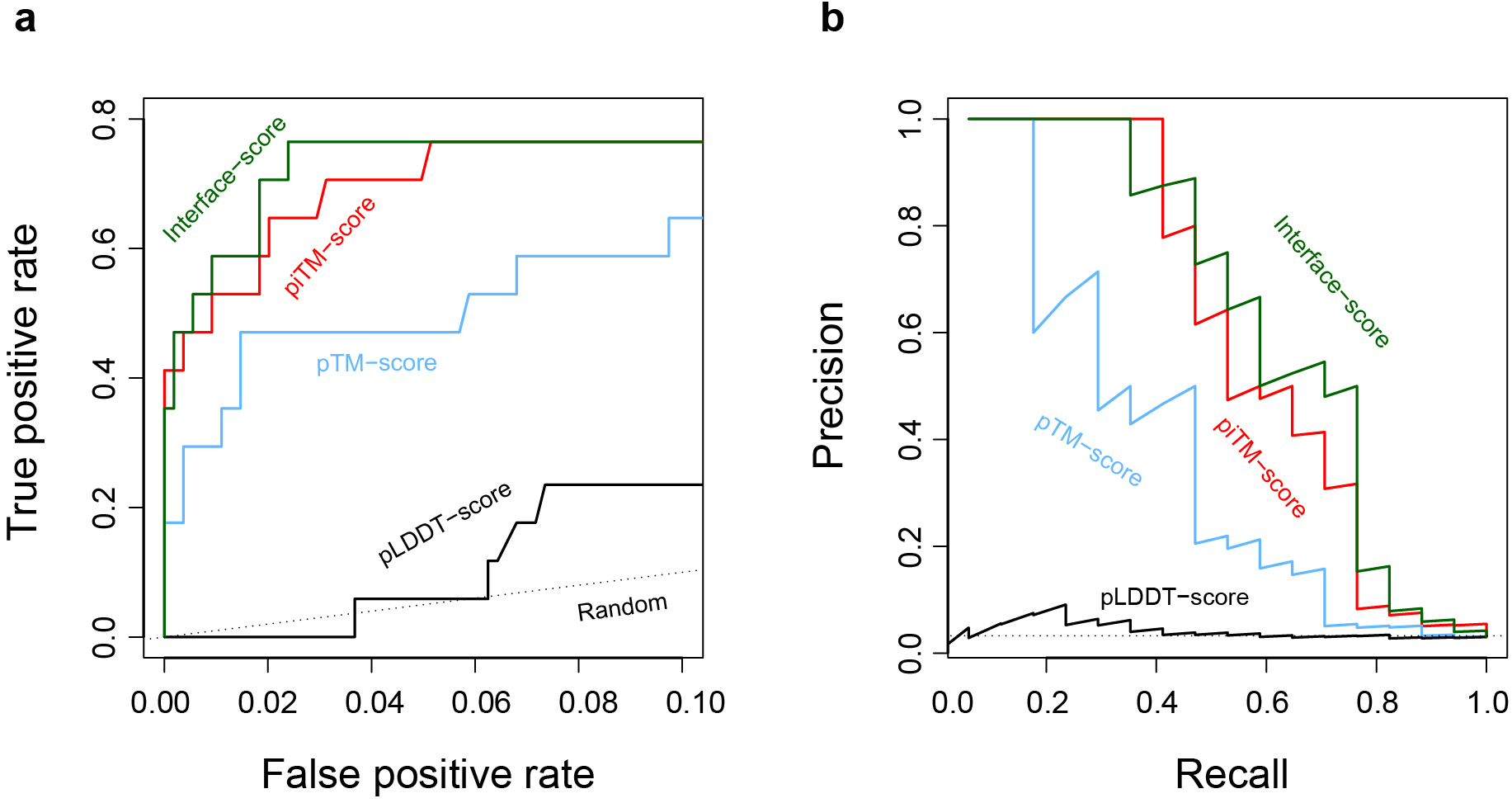
Identification of true interacting protein pairs in the all-against-all pool for the CP17 set by various confidence metrics. (**a**) Receiver operating characteristic curve and (**b**) the precision-recall curve. The random curve is the expected result by randomly guessing interacting protein pairs.

### Application to the *E. coli* proteome

The *E. coli* proteome consists of about 4,300 protein sequences. An all-against-all run with AF2Complex would require about 0.5 to 2.4 million computing node hours on the Summit supercomputer, which is beyond our allocation. Instead, we focused on a “gold standard” set of 701 PPIs previously curated largely from high-throughput experiments, and a set of 6,849 randomly selected, putatively non-interacting pairs (see Methods). Here, we have two goals: one is to test AF2Complex on a large-scale; the other is to build complex models for some known interacting protein pairs whose structures are difficult to determine experimentally. We speculated that the “gold standard” set contains pairs from a large assembly that are not necessarily in direct contact, e.g., as in ribosome. To test this hypothesis, we divided the positive set into subsets, whereby each monomer in the subset does not appear more than *C* times in the putative interacting set (the full set is covered when *C* < ∞). Fig. 5 shows the corresponding ROC and PR curves for these sets. The ROC curve displays a clear trend in which higher *C* values correspond to lower true positive rate or recall. This analysis suggests that some of the pairs in the positive set do not interact directly, yielding low or even zero scores. When we considered proteins that appear only once (i.e., *C* = 1), we obtained a result that largely recapitulates the benchmark performed above, with a slightly lower AUC_0.1_ of 0.60. The AUC_0.1_ drops to 0.50 at *C* = 3, 0.40 at *C* = 5, and 0.22 for the full set, likely due to the inclusion of more non-direct interacting pairs as *C* increases. Further analysis also found a barrier to accurate modeling is the lack of the exact context of the protein-protein interaction. For example, the chaperonin protein GroL is the most frequent monomer in this set, appears in 79 pairs. It is part of a large assembly that requires seven GroL copies forming a ring stacked with another heptameric GroE ring. However, when GroL was modeled alone with another putative partner, we only found models with low confidence scores. Despite this difficulty, the result suggests that AF2Complex performs as expected in a large-scale test. Of the positive set, among the predicted models with confident scores (interface-score > 0.45), we found that about 40% of these predictions have not been experimentally characterized (defined if both monomers share > 70% sequence identity with sequences found in the same PDB entry). Therefore, novel discoveries are expected from these models.

**Fig. 5.**
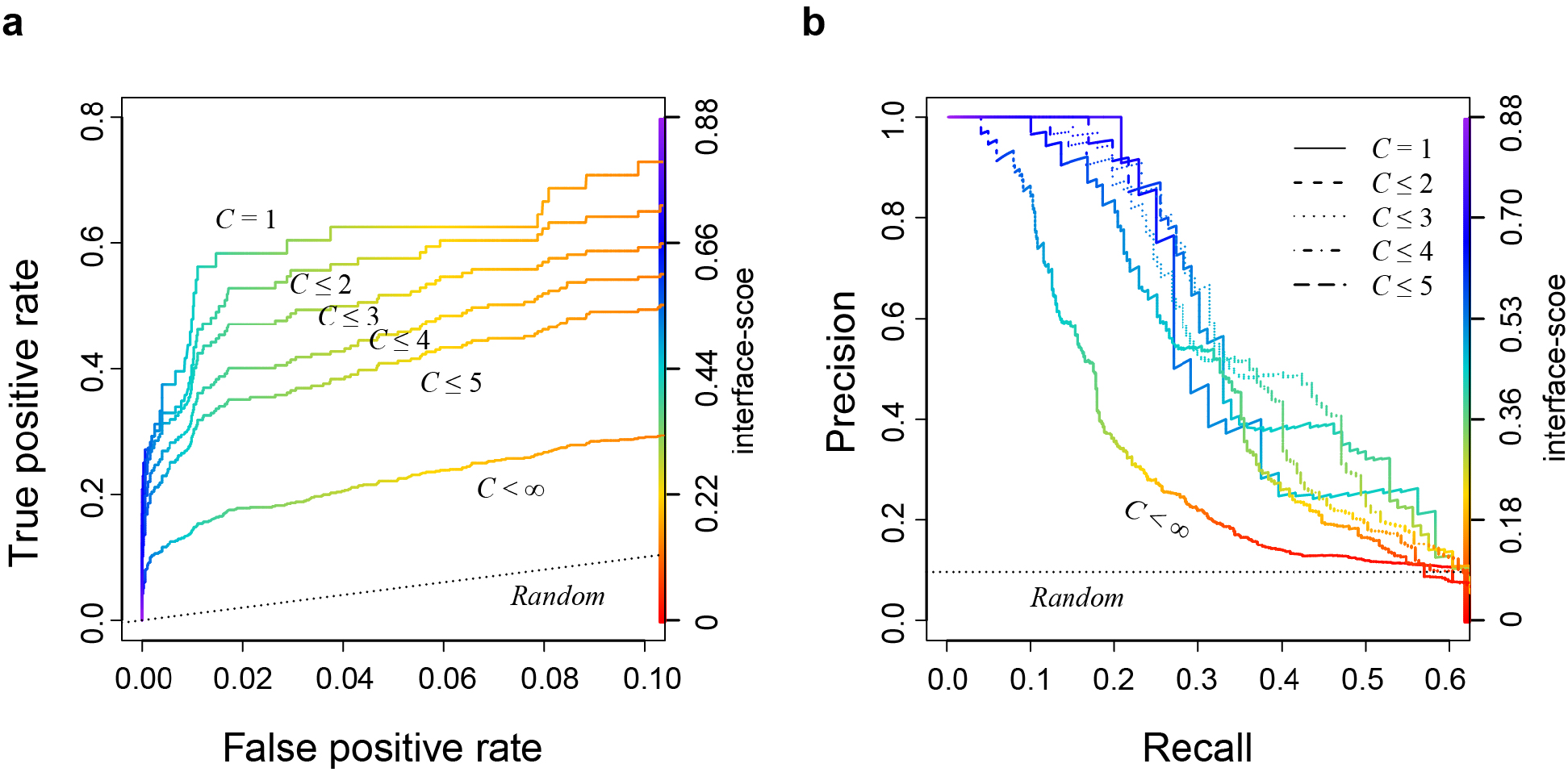
A Large-scale test on the *E. coli* proteome suggests that many pairs previously thought to interact directly are likely in assemblies of components that are not necessarily in direct contact. The interface-score was used as the varying metric to derive the (**a**) ROC curve and (**b**) the precision-recall curve. For a dimer target, *C* is defined by the maximum of the appearances of its two monomers in this data set.

### Structural models of cytochrome *c* biogenesis system I

Two *E. coli* targets with high-confidence models, CcmE/CcmF and CcmF/CcmH, caught our attention. These proteins belong to the cytochrome *c* maturation (Ccm) system I, which is composed of eight constituents (CcmABCDEFGH)[33, 34]. We note that *E. coli* CcmH has a fused C-terminal domain that appears as the standalone protein CcmI in other species with a similar Ccm system. As illustrated in Fig. 6a, it is thought that the Ccm system I consists of two modules: module 1 includes CcmABCD and is responsible for acquiring and loading a heme molecule onto the heme chaperone CcmE[44]. CcmE then shuttles the heme from module 1 to module 2, composed of CcmFGH, where the heme is delivered to CcmF[45]. Subsequently, CcmFGH covalently attaches the heme to nascent cytochrome *c*-type proteins[46, 47].

**Fig. 6.**
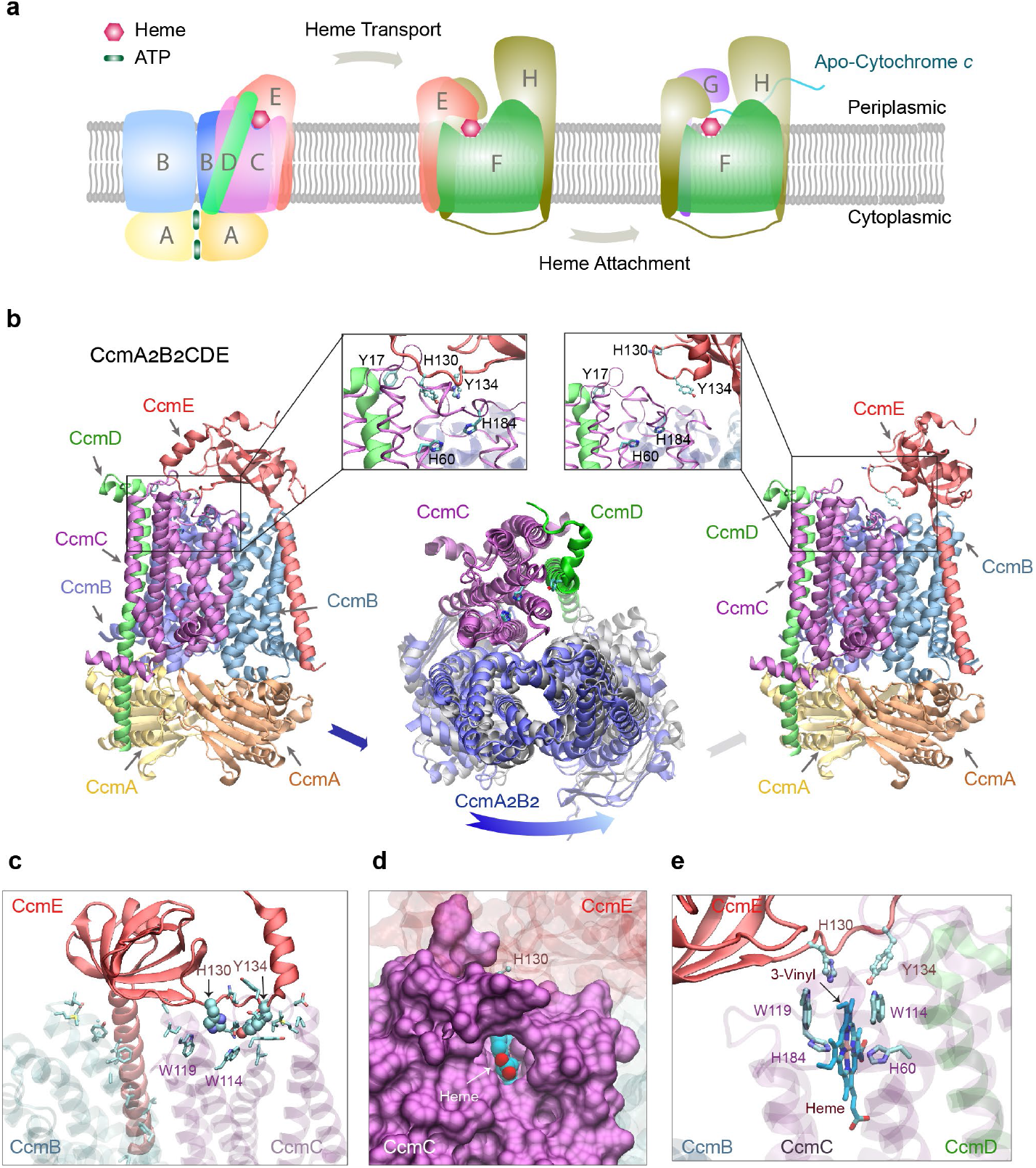
*E. coli*. cytochrome *c* maturation system I. (**a**) An illustration of the components and function of the Ccm I system, composed of eight proteins named CcmABCDEFGH. The system covalently attaches heme molecules to cytochrome c proteins via three functional complexes. (**b**) Two models (left and right panels) of one complex: CcmA_2_B_2_CD engage CcmE (left panel) and disengage CcmE (right panel). which loads a heme from CcmA_2_B_2_CD and chaperones it to CcmF. Insets show conserved residues implicated for heme binding in CcmC, CcmD and CcmE, respectively. Conformational differences between these two models are shown in the middle panel, where the backbone of CcmC was used to superimpose the two models. Viewed from the top (the periplasmic side), the two conformations of CcmA_2_B_2_ are displayed in blue and grey. Movement relative to CcmC is evident in CcmA_2_B_2_ but not in CcmD, which appears tightly coupled with CcmC. For clarity, CcmE is omitted in this superposition plot. (**c**) A view of interactions between CcmCD and CcmE in their engaged structural model shown in the left panel of (a). CcmCD representations are transparent for clarity. The sidechain of interacting residues (defined by heavy atom distance 4.5 Å) are shown. His130 and Try134 of CcmE are shown in the Van der Waals representation, and other interacting residues, including the Trp114 and Trp119 from the heme-binding WWD domain of CcmC, are shown in the licorice representation. (**d & e**) Views of a heme molecule docked into the putative binding-pocket in CcmC, using the structural model in which is CcmE bound to CcmA_2_B_2_CD as the initial apo-structure. A pore for heme access in CcmC manifests, where CcmC is shown in a surface representation (d). The heme is shown in van der Waals (d) and Licorice (e) representations. The vinyl group expected for His130^CcmE^ attachment is marked in (e).

Many mechanistic details of the Ccm system are still unclear, in part because there are no structures of the assembled modules. To date, the best effort is a partial model of the CcmCDE complex generated using co-evolutionary analysis[48]. The main reason for this knowledge gap is that the assemblies involve transient but essential interactions among membrane proteins (except for CcmA) that are difficult to capture experimentally. We sought to address this knowledge gap with AF2Complex by modeling about two dozen combinations of Ccm components (see Methods). From these computational experiments, we present the confident models of three assemblies, A_2_B_2_CDE, EFH, and FGH, the last one with and without apocyt peptides, all with high interface-score (0.82, 0.56, and ~0.72, respectively). These models are likely biologically relevant, and we were able to model the heme in the expected sites in all these models.

#### CcmA_2_B_2_CDE

First, we identified two different conformational states in the top two highest-ranking models of A_2_B_2_CDE (Fig. 6b). Extensive interactions are observed between CcmCD and CcmE in one model (Fig. 6b left panel), where the heme chaperone CcmE extends its heme-binding loop, characterized by two conserved residues His130^CcmE^ and Tyr134^CcmE^, to reach the heme binding site (HBS) in CcmC that includes His60^CcmC^ and His184^CcmC^, and another conserved residue, Tyr17 of CcmD. The heme-binding loop of CcmE is oriented away from the HBS of CcmCD in the other model (Fig. 6b right panel), in which the interactions between CcmC and E disappear, and the only remaining inter-protein residue-residue contacts are between a pair of transmembrane helices of CcmC and E. This conformation presumably corresponds to a resting state before heme loading, after heme unloading, or both. The disengagement of CcmE leads to conformational changes within CcmA_2_B_2_CD (Fig. 6b center panel). Using CcmC as the reference to superimpose the two complexes, movement is evident in CcmA_2_B_2_ but not in CcmD, which appears tightly coupled with CcmC. The root mean squared deviation (RMSD) of the CcmC backbone is ~1 Å. Corresponding to this movement, the number of residue-residue contacts between CcmB and CcmC drops by 45%, and the CcmBC protein-protein interfaces display larger changes between the two models than the other interfaces (similarity score, IS-score, of 0.65 versus scores >0.84). These large conformational changes could be the result of ATP hydrolysis within CcmA. As previously proposed[44], the energy of the hydrolysis could be harnessed to release the cargo-loaded CcmE.

Extensive contacts are present in the model with CcmC tightly bound with CcmE. The interactions involve Trp114 and Trp119 of a tryptophan-rich loop (WWD domain[49]), sitting at one edge of the binding pocket of the heme (Fig. 6c). A pore is visible between transmembrane helices 2 and 5 of CcmC and exposes a heme molecule bound to CcmC (Fig. 6d). The pore may allow the access of heme from the outer leaflet, but no channel for potential heme trafficking is present within CcmC as previously speculated[49]. The role of His60^CcmC^ and His184^CcmC^, predicted heme iron-coordinating residues, is confirmed as well (Fig. 6e). His130^CcmE^ is only ~4 Å away from the 3-vinyl of heme (IUPAC numbering), which is the proposed site of covalent attachment to His130^CcmE^ to complete the heme delivery to CcmF.

#### CcmEFH

Next, we addressed the question of how a heme-carrying CcmE could deliver heme to the CcmFGH complex. We obtained a confident model in which CcmE is in complex with CcmF and CcmH (Fig. 7a). CcmE interacts with CcmF such that the HBS of CcmF faces the HBS of CcmE, which has a heme-handling motif like that in CcmC. The distance between His130^CcmE^ and His303^CcmF^ is ~6 Å (Fig. 7a inset). CcmF has two heme-binding sites: one (P-heme) for the cytochrome attachment and the other accessory heme (TM-heme) that assists the ejection of the P-heme[34]. His303 is highly conserved and is known to coordinate the Fe cation bound to P-heme. Similar interacting poses between CcmE and CcmF were obtained in models generated in the absence of CcmH.

**Fig. 7.**
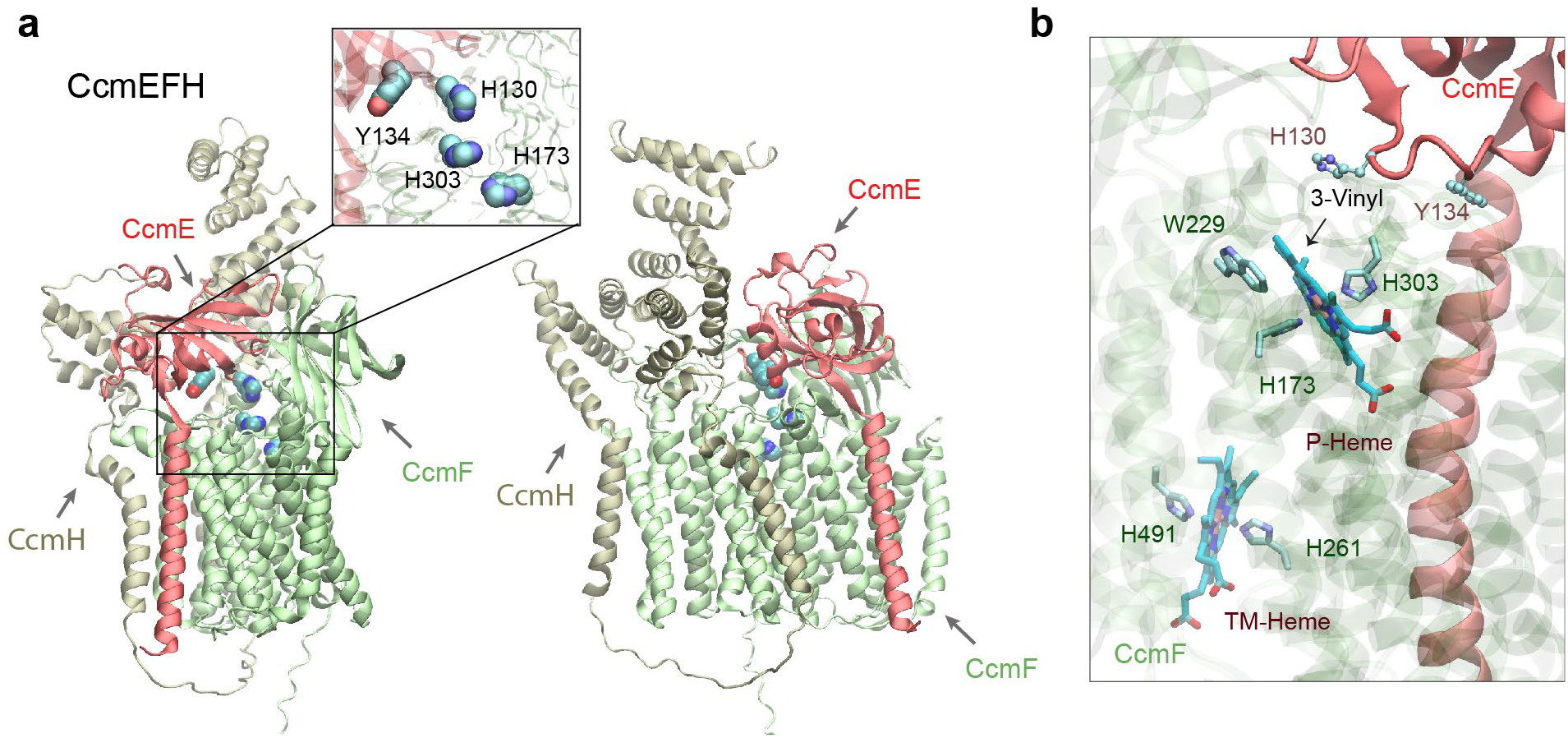
Structural models of the CcmEFH complex from *E. coli*. (**a**) CcmE is believed to deliver a heme to CcmF. Two views of a top model generated by AF2Complex are shown in the cartoon representation. The inset shows the key heme handling residues, His130 and Tyr134 of CcmE and the two histidines of CcmF. (**b**) Two heme molecules computationally docked to expected heme-binding sites of CcmF using the model shown in (a). The critical heme iron coordinating residues, His173 (P-His1) and His303 (P-His2) for the P-heme delivered by CcmE and eventually attached to an apo-cytochrome c protein and His261 (TM-His1) and His461 (TM-His2) for the co-factor TM-heme are also shown.

Consistently, the heme-bound model displays the P-heme coordination role of His303^CcmF^ and His173^CcmF^, and the TM-heme is coordinated by His261^CcmF^ and His461^CcmF^ (Fig. 7b). TM-heme was co-crystallized with CcmF from *Thermus thermophilus* (*Tt*CcmF) in a recent published X-ray structure[50]. Without using this structure as a template, our predicted CcmF model has a TM-score of 0.92 compared to the experimental structure. However, a critical loop containing the equivalent histidine of *E. coli* His303^CcmF^ is missing in the *Tt*CcmF structure, suggesting that the P-heme binding pocket might be flexible. Indeed, only one of the two expected tryptophans (W229 and W236) from the WWD domain is in contact with the P-heme in the heme-bound model. Moreover, CcmF in our model does not expose a pore as observed the *TtCcmF* structure, in which the pore was speculated to enable P-heme access[50]. The structural difference may be attributed to lipid molecules occupying the pore in the crystal structure but absent in our computational model.

#### CcmFGH

After the heme is delivered to CcmF, the final step performed by this system is the attachment of the heme to apocytochrome *c* (apocyt *c*). This step involves a complicated mechanism that is not fully understood[47]. However, our model of the CcmFGH complex provides structural insights into the mechanism (Fig. 8a). First, we note the mobility of the N-terminus of CcmH (which would be the full CcmH in many other systems that also have CcmI but is fused to CcmH in *E. coli*). In the absence of CcmE, the N-terminus of CcmH occupies the site otherwise occupied by CcmE, essentially moving closer to the HBS of CcmF. This configuration leaves an opening for CcmG, another thiol-disulfide oxidoreductase (like CcmH), now sitting at the site previously occupied by the CcmH N-terminal domain. Remarkably, the CcmFGH complex is arranged such that a reaction groove is formed, in which an apocyt *c* can be sequentially passed among the CXXC motifs of CcmG (Cys80 and Cys83) and CcmH (Cys43 and Cys46) to reach the HBS of CcmE (Fig. 8a).

**Fig. 8.**
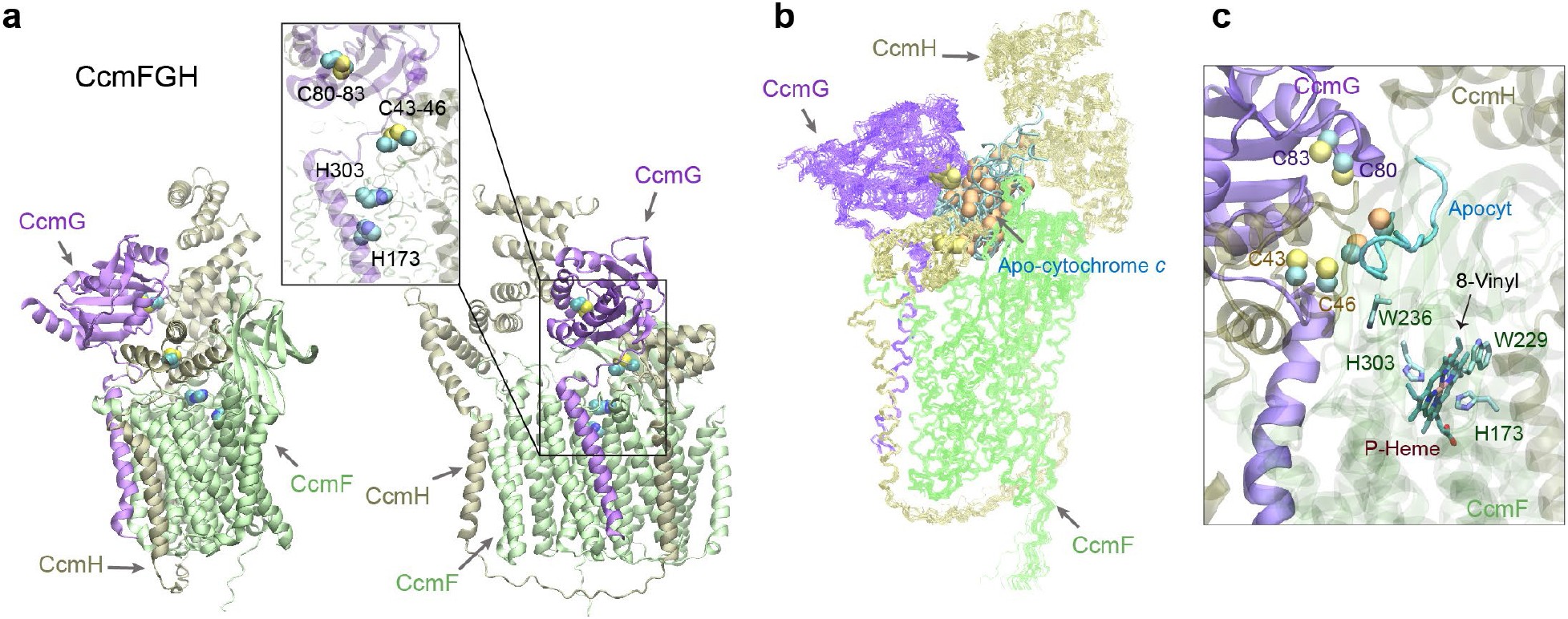
Structural models of the CcmFGH complex with and without apo-cytochrome c substrates from *E. coli*. (**a**) Two views of a top model of CcmFGH are shown in the same orientations as the two views in Fig. 7a, respectively. The CcmH N-terminal domain moves closer to the heme-binding sites of CcmF, leaving space to accommodate CcmG that now binds CcmF with CcmH. Critical cysteines of the CXXC motifs of CcmG (Cys80 and Cys83) and H (Cys 43 and Cys46), and the P-heme-binding histidines (His 173 and His303) of CcmE are shown in the vdW representations in the inset. (**b**) Superimposed 22 structural models of the CcmEFH and apocyt acceptors by AF2Complex. CcmFGH are shown in lines, and apocyts are shown in cyan tubes. All apocyts are found within the same groove formed by the three Ccm proteins. The superposition uses the backbone atoms of CcmFGH as the reference. The sulfur atoms from the CXXC motifs of apocyts are shown in orange spheres to differentiate from those of CcmGH. (**c**) A heme molecule computationally docked to the P-heme’s binding site in CcmF using one of the models shown in (b). One of the Cys residues of apocyt is found within 4 Å from Cys46 of CcmH. The distance between the other Cys residue of apocyt and the 8-vinyl group of the heme is about 16 Å.

To investigate further, we modeled CcmFGH together with 11 different apocyt peptides, each containing one or two CXXCH motifs (see Methods). Remarkably, in all top models, the apocyts are invariably located in the predicted reaction groove (Fig. 8b). Physical contact between Cys46^CcmH^ and an apocyt cysteine is present in some models (Fig. 8c). We further modeled a heme in the expected P-heme site of CcmF. The heme-bound model is largely as expected, but more conformational changes are necessary for apocyt-heme attachment, as the cysteines of the apocyt are still ~16 Å away from the 8-vinyl group of the P-heme. The speculation echoes large conformational changes upon heme-binding revealed in recently determined structures of CcsBA, a cytochrome *c* synthetase from a different Ccm system[51].

Of the previously proposed mechanisms[47], our model supports the following: after entering the reaction groove, apocyt *c* is first reduced by Cys80 and Cys83 of CcmG. Next, the reduced apocyt attacks one of Cys43 and Cys46 of CcmH to form a mixed disulfide. This intermediate complex then retrieves the heme acquired by CcmF, and subsequently the mixed disulfide is resolved by the second cysteine of CcmH. Finally, the holo-cytochrome *c* is released. The CXXC motif of CcmH then reverts to the oxidized state, and CcmG dissociates to be reduced by the thiol-disulfide interchange protein DsbD. In all models of CcmFGH and also models from separate modeling of a CcmGH complex, the CXXC motifs of CcmG and H are separated by ~15 Å, and hence the reduction of a mixed disulfide between these two motifs as proposed in an alternate mechanism[47] is unlikely according to these models. Interestingly, the two CcmH domains, encoded in two ORFs in some organisms but fused together in *E. coli*, are linked by a long loop without direct interactions. This is an exception to the notion that fused proteins directly interact[4]. In this case, both CcmH domains instead interact with a third protein, CcmE. The function of the CcmH C-terminal domain remains unclear, but likely involves interactions with an apocyt *c*.

## Discussion

Our findings clearly demonstrate that AF2 can be adapted to predict the structure of protein complexes at much higher accuracy than classical docking approaches, even if the docking approaches use monomeric structures predicted by AF2. One reason is that predicting all the protein structures involved in a complex simultaneously by AF2Complex may overcome issues associated with rigid-body docking. Importantly, we have shown in multiple benchmark tests that high-quality complex prediction can be achieved *without* using paired MSAs as input, which also significantly enhances AF-Multimer and returns more physical complex models. This feature may lower the barrier for applications including some challenging cases in which pairing MSAs is impractical.

But why is it possible to achieve successful complex modeling without using paired MSAs? After all, for predicting single chain structures and also for predicting complex structures, MSAs for each individual protein are still necessary and important. One reason is that sophisticated deep learning models reduces their reliance on large MSAs[1, 52]. Moreover, we speculate that the accurate amino acids packing capabilities offered by AF2 deep learning models may be another key reason[1, 3]. In particular, the structural module of AF2 ignores the sequential order of amino acids and has likely learned energetically favorable patterns among packed amino acids. If these patterns are applicable universally to amino acids of either intra- or inter-proteins, success is then expected. Empirically, the results above indicate this is likely the case. After all, protein-protein interactions are not physicochemically different from what drives protein folding in the first place. Their interface structures likely have been learned during the training of AF2 deep learning models for monomeric protein model prediction.

Furthermore, by assessing the confidence of a predicted complex model with carefully designed metrics, one may generalize this deep learning approach to predict direct protein-protein interactions. We demonstrate that the interface-score or piTM metric can be applied to effectively predict interacting pairs. When applied to an *E. coli* proteome, we were able to infer that some of the previously selected interacting pairs in the “gold standard” set are likely from large complex assemblies without direct interactions.

Using the *E. coli* cytochrome maturation system I as an example, we demonstrated that this powerful computation tool can be applied to interrogate a molecular system comprised of many proteins arranged in different packing orientations. By way of example, AF2Complex can generate highly confident models that depict the complexes involved in the loading, release, and delivery of a heme-chaperone, including a reaction groove responsible for the final attachment of a heme to an apo-cytochrome protein. Remarkably, high-confidence models were obtained for these assemblies that include multiple conformational states involving transient interactions with the heme chaperone and apocytochrome *c* peptides. Although the approach is currently limited to structural models without directly incorporating a heme, docking studies suggest that our models are consistent with known biochemical evidence, though other conformations are also expected.

One major hurdle to this bottom-up approach for predicting protein interactions is that the context of such a hypothetical complex is often unavailable a priori. For example, if a complex involves one homodimer and another monomer, it would be difficult to model if we only consider a single heterodimer. Another challenge is post-translational modifications. For instance, proper modeling of CcmEFH and CcmFHI requires the cleavage of the N-terminal signal peptide of CcmH to obtain biologically accurate models. Nevertheless, the power of a deep learning-based approach for predicting direct protein-protein interactions has been demonstrated. It is expected to contribute profound structural insights into the understanding of many biological molecular systems.

## Methods

### AF2Complex workflow

AF2Complex was initially built upon the official release of the source code and monomer neural network models of AlphaFold2 (version 2.0.1)[1], and subsequently upgraded to support the multimer neural network models of AlphaFold-Multimer (AF version 2.1.1). For the purpose of large-scale applications, the original data pipeline was separated from the neural network inference. We refer to the data pipeline and neural network inference portions as stage 1 and 2, respectively. The split allows us to derive input features for individual protein sequences and then reuse them to assemble input features for subsequent complex predictions. We used different sets of sequence libraries[53–56] and Protein Data Bank (PDB)[57] releases to generate appropriate input features for different test sets, as described below.

To generate the MSAs for predicting complex structures made of *N* distinct protein sequences, each with a length *L_i_* and a stoichiometry number *S_i_* (*i* = 1… *N*), we apply Algorithm 1 to the MSAs of individual proteins. The application creates a new set of complex MSAs, whose length is the sum of all individual sequences including multiple copies in the case of homo-oligomers, and whose depth is the sum of the depths of all individual MSAs. The complex MSAs are primarily composed of gaps, except for the regions in which each individual target sequence has its own window of MSAs (see Fig. 1 of the Main text for a schematic example).

#### Algorithm 1 Complex MSA Creation

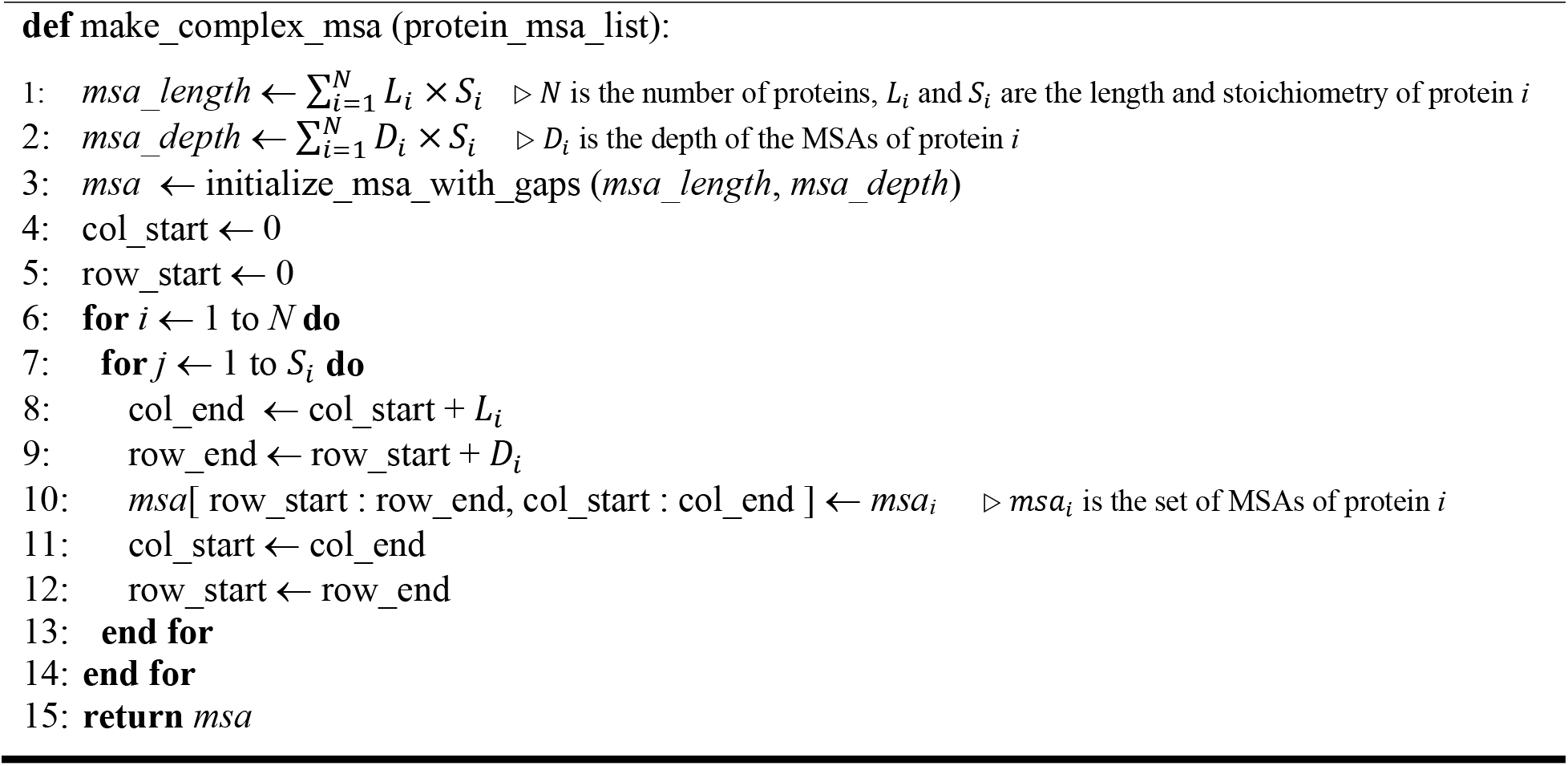

When supplied to a monomer AF DL model, the “*residue_index*” input feature for the target complex was modified by increasing the residue indices of individual protein sequences by *b*(*p* – 1), where *b* is an arbitrarily chosen number of 200, which satisfies the condition larger than the coverage of the relative positional encoding at 32 (that is, the sequential distance between two residue indices |*i* – *j*| ≤ 32, see Algorithm 4 of the Supplementary Method of reference[1]), and *p* denotes the index of each monomer starting from 1. Likewise, the template structures for individual proteins were also collected for the complex prediction. However, we did not specifically search for and supply a complex template for the target. Without a note, the neural network models used for subsequent structure prediction were the five monomer AF2 models, each with a fine-tuned head for predicting paired alignment errors, which allows the prediction of the TM-score (pTM) [1]. We took advantage of this head for deriving metrics for evaluating complex predictions (see below). In the tests described in this study, we increased the number of recycles up to 20, depending on sequence lengths; the maximum number of recycles for target sequences longer than 500 residues was progressively decreased to reduce computational costs. The recycle steps were stopped early if the backbone *C_α_* distogram converged [20]. In practice, these options may be adjusted by end users.

### Metrics for complexation evaluation

Previously, we introduced the interface TM-score (iTM-score) and interface-similarity score (IS-score) for measuring the structural similarity between protein-protein interfaces[31]. Both scores were introduced to deal with issues associated with the TM-score, which is not ideal for comparing structure similarity of protein complexes[31, 58]. In this study, we used a similar concept but modified it accordingly for estimating the confidence of a predicted complex model. We first introduce the predicted interface TM-score, piTM,

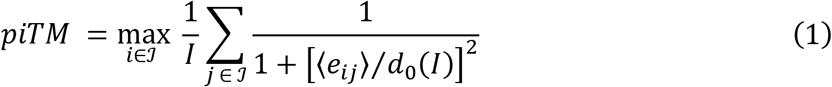

where ℐ is the set of interface residues observed in the predicted model structure, and the cardinality of ℐ is the total number of interface residues *I* ≡ |ℐ|. Using the local reference frames of interface residue *i*, the predicted alignment error head of AF2 gives an estimated distance ⟨e_ij_⟩ for interface residue *j* from its position in the experimental structure[1]. The piTM score is the optimal rotation/translation that gives the best estimated score, and d_0_(*I*) is a normalization factor given by,

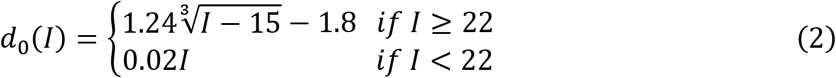

Note that we adjust the original formula of *d_0_* to better deal with the cases where a low number of contacts are observed. Furthermore, we define the interface-score *S* as the follows,

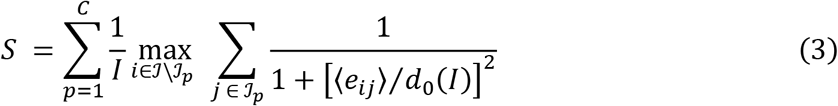

which is similar to piTM, but we now calculate a piTM score for each protein chain *p* of the complex separately and then sum the scores. Each chain *p* has an observed number of interface residues ℐ_p_, and ℐ is the union of ℐ_p_. The optimal local reference for calculating the score for chain *p* can only be selected by interface residues not belonging to chain *p*. An important difference between our metrics and the ipTM score introduced in ref [35] is that we focus on interface residues (versus full chains[35]), which is the most relevant for our interaction predictions.

### CASP14 multimeric targets

We modeled all assembly targets if the total size of an assembly is less than 3,000 residues, which is a limit imposed by our available computing resources. The sequence libraries employed for our predictions are UniRef90 created in 2020-01[59], and the reduced BFD, MGnify, and Uniclust30 libraries[53–56] provided with the AF2 release. All these libraries are composed of sequences available prior to CASP14. For template retrieval from the PDB, we restricted ourselves to structures released before the starting date of CASP14 as well. From the final predictions, we selected some difficult targets with promising results to present.

### CP17 set

This set was taken from ref [22] and consists of 17 heterodimer targets released after 2018-04-30, on which the experimental structures were collected from the PDB for training the AF2 neural network models. We used the same sequence library as above but restricted the structural templates from the PDB to those dated before the same cutoff date for the AF2 training structures. In the benchmark test of the 17 targets, three independent runs were carried out, and each generated five top models. We evaluate the overall top-ranked model of the 15 models according to the interface-score. This protocol was used to evaluate both monomer and multimer AF neural network models. In the second benchmark test of identifying the 17 heterodimers out of all 561 pairs of these 34 individual protein sequences of the dimers, one single run was conducted for all pairs. Up to 20 recycles were allowed in each of these runs.

The top-ranked (by the interface-score) models were compared with their corresponding experimental structures with the programs IS-score[41] and DockQ[43]. Inaccurate, acceptable, medium, and high-quality models are defined by DockQ score regimes [0.0, 0.23), [0.23, 0.48), [0.48, 0.80), and [0.80, 1.0], respectively.

### Dimer1193 and Oligomer562 sets

To create benchmark sets for objective evaluations of the AF2 models, we curated two sets from the experimental structures released recently in the PDB. First, we retrieved the information of all 120,703 protein assemblies on Jan 7, 2022 from the PDB. They were split into two subsets, 92,047 complex structures (Earlier Set) released prior to Apr 30, 2018, and 24,853 assemblies (Later Set) released after that date, which is the cutoff date for the structure collection used to train the AlphaFold DL models including both monomer and multimer DL models[35]. We assume all assemblies in the Earlier Set were used for DL training and removed all the “easy” homologs in the Later Set by using the 30% sequence identity clusters of all protein sequences provided by the PDB. For a complex in the Later Set to be considered further, at least one protein chain of this complex must not be found within the same 30% sequence identity cluster as any chain of any structure in the Earlier Set. This selection procedure left 8,544 assembly structures. To further remove the redundancy among them, we applied the same sequence identity criterion and arbitrarily selected 3,353 structures that have at least one chain not found in the same 30% cluster as any chain in any other complex structure within these 8,544 structures. A maximum length of 1,480 residues was also applied during the selection to the total size of the oligomer (i.e., the residue count of all individual chains) to prevent GPU memory overflow, a limitation imposed by the memory capability of the GPUs available for our tests.

The sequences and coordinates of the 3,353 assemblies were subsequently examined. We further require that, in a good target, each protein chain shares a protein-protein interface of at least 20 residues with another chain of the complex. This requirement removed many complexes with short peptides and left 1,884 assembly structures, of which 47 entries with non-standard amino acids in their PDB SEQRES records were removed, as we used the sequences given by SEQRES as the input sequences to model prediction. The remaining 1,838 assemblies consist of 1,275 dimers and 563 higher order oligomers. One entry in the oligomers failed to generate the input during our evaluation of AF-Multimer, and we removed it from further evaluation. These two final sets are called Dimer1275 and Oligomer562, respectively. Because the mapping of chains between model and experimental structure is an open issue for automated assessment of high-order oligomers, primarily due to the combinatorial growth of identical or homologous copies, we mainly focused on the dimer set in this study. However, we provide an evaluation of the physicality of the predicted Oligomer562 models.

In addition, AF-Multimer uses different template identification protocols between the multimer and monomer predictions. The multimer data pipeline uses a protocol that searches essentially all PDB structures up to an optional cutoff date (Apr. 30, 2018, in our benchmark scenario), but the monomer protocol searches only a subset of PDB structures (PDB70, representing single chains at 70% sequence identity cutoff). All targets of Dimer1275 have identified four templates in the multimer protocol, but 82 targets have none or fewer than four templates identified in the monomer protocol that AF2Complex relies on for input generation. To alleviate this unfairness, we removed these 82 dimers, leading to the final Dimer1193 set for the benchmark tests. The union of Dimer1193 and Oligomer562 are named as Oligomer1755.

### *E. coli* sets

The sequence of *E. coli* strain K12 was downloaded from UniProt[59] (Proteome ID UP000000625). We used a positive and negative protein-protein interaction set curated previously[27, 32]. We first filtered out pairs whose total size is longer than 1,480 residues, which is a limit imposed by the 16 GB GPU memory per node on the Summit supercomputer. Filtering led to 701 pairs from the positive set. Because the original negative set is too large to run all, we randomly selected 6,849 pairs from them, which yielded roughly a 1:10 ratio between the positive and negative set.

### *E. coli* CcmI system modeling

Three rounds of modeling were performed for this system composed of eight proteins, CcmA, B, C, D, E, F, G and H. In the first round, based on the literature, we tested 22 combinations of these proteins. Among them, top models with high confidence interface-scores and literature corroborations were presented for three assemblies, CcmA_2_B_2_CDE, CcmEFH, and CcmFGH. All models were generated with the AF2 monomer DL models by applying up to 20 recycles. Application of the AF-Multimer DL models with AF2Complex did not yield better models with higher confident scores and are thus not presented.

In the second round, we tested the interactions between various apocyt *c* peptides and the CcmFGH complex. A cytochrome *c* protein, NrfA of *E. coli*, was arbitrarily chosen for modeling. NrfA contains five CXXC motifs for covalent heme attachment[60]. We cropped peptides spanning the heme binding motifs, including 8 peptides with one CXXC motif and three with two motifs. The lengths of these apocyt *c* peptides range from 8 to 52 residues, mostly around 18 AAs. During modeling, an apocyt *c* substrate and CcmFGH were folded simultaneously by the DL models, in contrast to typical docking, whereby a substrate is placed into a putative binding site of a folded protein or complex structure. By simultaneously modeling both the receptor and the acceptor, one might obtain a better complex structure that requires large conformational changes due to interactions. Note that we did not re-generate the MSAs for each peptide substrate. Rather, we cropped out the input features for each peptide from the input features, including the MSAs, of the full NrfA sequence. A domain cropping option implemented in AF2Complex enables this practice, which is convenient and likely more accurate in comparison to regenerating MSAs using partial sequences, especially for short peptides.

In the last round, we modeled heme *b* molecules in their putative binding sites of CcmC or CcmF of top models. Because AF2 does not currently support the incorporation of cofactors and other prosthetic groups into structural models, we ran the Rosetta *relax* application[61] using the ‘*-in:auto_setup_metals’* option to model heme-bound systems starting from the AF2C models. The *molfile_to_params.py* script was used to generate the required parameters for heme *b*. In some cases, the positions of histidine residues were adjusted to place them in proper position for axial coordination prior to Rosetta refinement.

### Performance evaluation

Standard metrics were applied to the benchmark tests on the CP17 and *E. coli* sets, both consisting of a true positive and negative set. The predictions were labeled using the pre-defined classification and the numbers of true positives, false positives, true negatives, and false negatives were then designated as TP, FP, TN, and FN, respectively. Performance measures are defined as follows,

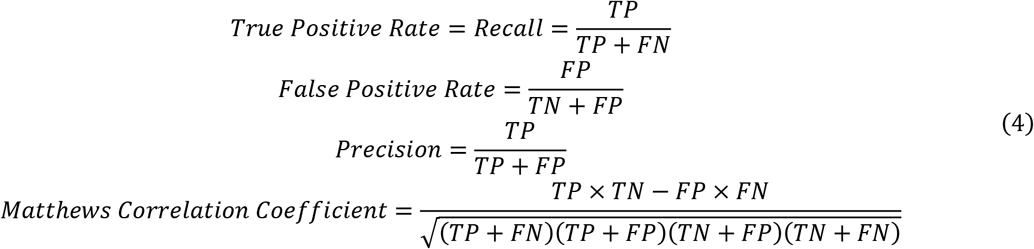

We also employed the normalized AUC_0.1_, which is the area under the ROC curve up to an FPR of 0.1, divided by 0.1. ROC curves were plotted using ROCR [62].

### Interface clash indicator

To characterize the clashes observed in the predicted computational models, we defined an interface clash indicator *χ* ≡ *N_iRes_/N_iCnt_*, where *N_iRes_* is the number of interface residues, and *N_iCnt_* is the number of interface residue-residue contacts. An interface residue-residue contact is defined if any heavy-atom of a residue of one chain is within 4.5 Å of any heavy-atom of a residue from another chain. The value of χ ranges from 0 to 2 with values close to zero indicating a significant number of clashes. For experimental structures, χ is typically between 0.6 and 1.2. The metric can be directly applied to un-relaxed models to detect severe clashes. In practice, this evaluation can help to save computing time on unphysical models that are unlikely to be fixed by the relaxation protocol of AF2.

### Statistical tests

To test the hypothesis that the model quality of one model is better than another one in terms of a scoring metric, e.g., the DockQ score, we used the Wilcoxon signed-rank test, a non-parametric statistical test, because the score distributions do not follow normal distributions. The tests were paired, as every method makes a prediction on the same set of targets. All tests are one-tailed.

### Computational costs

The development tests and predictions on CASP14 assembly targets were carried out locally using about 10 workstations each with four Nvidia RTX6000 GPUs, where each GPU has 24 GB memory. The benchmark tests on CP17 and *E. coli* sets were performed on the Summit supercomputer at Oak Ridge National Laboratory. A Singularity container was built to run AF2 on Summit[63, 64]. An AF2Complex run of ~7,000 pairs of proteins using 923 nodes required about 2 hours in wall clock time. Each node has 6 Nvidia 16 GB V100 GPUs. For an individual target of fewer than 1000 residues, models may be obtained within 20 minutes for each deep learning model using “*super*” mode, which is a preset of configurations used with AF2Complex for this study.

## Data availability

The source code of AF2Complex is available at https://github.com/FreshAirTonight/af2complex. Benchmark data sets of CP17, Dimer1193, Oligomer562, and the full *E. coli* proteome, including pre-generated input features to AF2Complex, and the top computational models of *E. coli* Ccm system I are available at Zenodo https://doi.org/10.5281/zenodo.6084186.

## Acknowledgments

We thank Ada Sedova for coordinating the deployment of AlphaFold2 on Summit at Oak Ridge and critical reading of the manuscript, Ryan Prout, Subil Abraham, Wael Elwasif, N. Quentin Haas for building a Singularity container, and Mark Coletti for providing Dask scripts for running AF2. We thank Jessica Forness for proofreading the manuscript. This work was supported in part by the DOE Office of Science, Office of Biological and Environmental Research (DOE DE-SC0021303) and the Division of General Medical Sciences of the National Institute Health (NIH R35GM118039). The research used resources supported in part by the Director’s Discretion Project at the Oak Ridge Leadership Computing Facility, and the Advanced Scientific Computing Research (ASCR) Leadership Computing Challenge (ALCC) program. We also acknowledge the computing resources provided by the Partnership for an Advanced Computing Environment (PACE) at the Georgia Institute of Technology.

## Author contributions

MG and JS designed the research, MG and DA wrote the source code, MG performed research and analyzed the data, MG and JP analyze the models of the *E. coli* Ccm system I, MG prepared the first draft of the manuscript, MG, JP and JS revised the manuscript, and all authors proofread the manuscript.

## Competing interests

Authors declare no competing interests.

## Supplementary Information

### Supplementary Text

After we finalized the first version (1.0) of AF2Complex, DeepMind released AF-Multimer (AlphaFold version 2.1.1)[1]. To assess and take advantage of the new deep learning (DL) models trained for predicting multimers, we adapted AF2Complex (version 1.2) to further support AF’s new multimer DL models with either paired MSAs or unpaired MSAs as the input. In the case of paired MSAs, they were generated using DeepMind’s data processing pipeline. However, in the model inference stage AF2Complex provides options to tweak runs according to users’ needs, such as unpaired MSAs and additional recycles rather than the default three recycles. We use “monomer” and “multimer” to differentiate these two sets of DL models of AF2 released in version 1.0.1 and 2.1.1, respectively.

Using CASP14 target H1060v4 as an example, we show both the strengths and weaknesses of the multimer DL models. As shown in Fig. 2d of the main text, using an AF2 monomer model, AF2Complex predicts an elliptical model, rather than the expected C12 symmetric model[2]. Using the multimer models released with AF-Multimer, we obtained a complex model with C12 symmetry, displayed in Fig. S1a. The C12 symmetric model has very similar dimeric interfaces as those observed in the monomer elliptical model (Fig. S1b), but the cyclic symmetry is broken in the previous model and is maintained only in the new model. Moreover, only one of the five multimer DL models generated such a model by using unpaired MSAs. It appears that using unpaired MSAs may be the key for generating this good model, because runs with paired MSAs by following the AF-Multimer workflow return unphysical models with severe clashes, a phenomenon akin to “chain collapse” observed previously[3]. Paired MSAs may have contributed to the issue. We shall revisit this topic below.

**Fig. S1.**
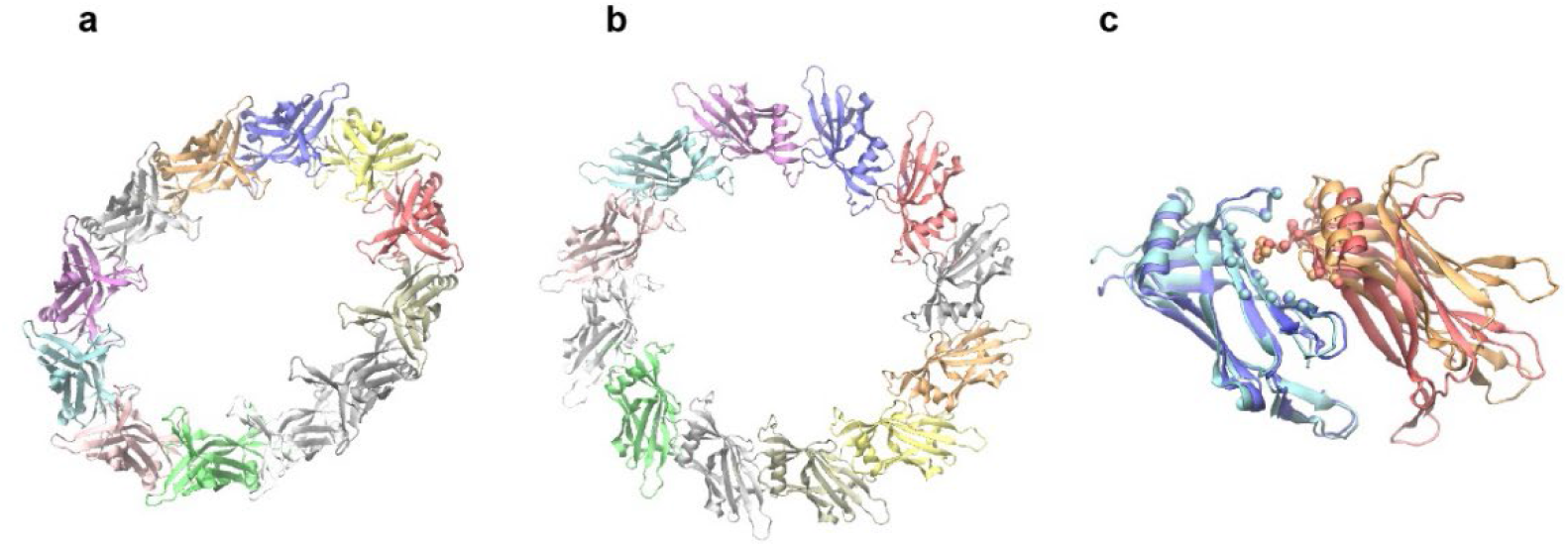
Top AF2Complex model of CASP14 target H1060v4. Structural model of AF2Complex generated with (**a**) an AF2 monomer DL model and (**b**) an AF-Multimer DL model. But are obtained with unpaired MSAs. (**c**) The superimposition of one pairs of monomers from this model and the top model obtained with an AF2 monomer DL model. The two copies of the proteins are colored cyan/orange (multimer model) and blue/red (monomer model). Backbones are shown in the cartoon representation and the Cα atoms of aligned interface residues are shown as spheres. The superposition was carried out with the program iAlign [4].

### Improvement with AF-Multimer’s deep learning models

To provide a comprehensive benchmark test with AF-Multimer’s DL models, we curated the Dimer1193 data set of 440 heterodimers and 753 homodimers from the PDB experimental structures that were released after Apr 30, 2018, the cutoff date of experimental structures used by AF-Multimer DL model training (see Methods). In addition, each dimer has at least one protein chain that shares less than 30% sequence identity with any chain found in any assembly structure released before the cutoff date in the PDB. This requirement removes “easy” cases where the DL models already learned a target during their training sessions.

**Fig. S2.**
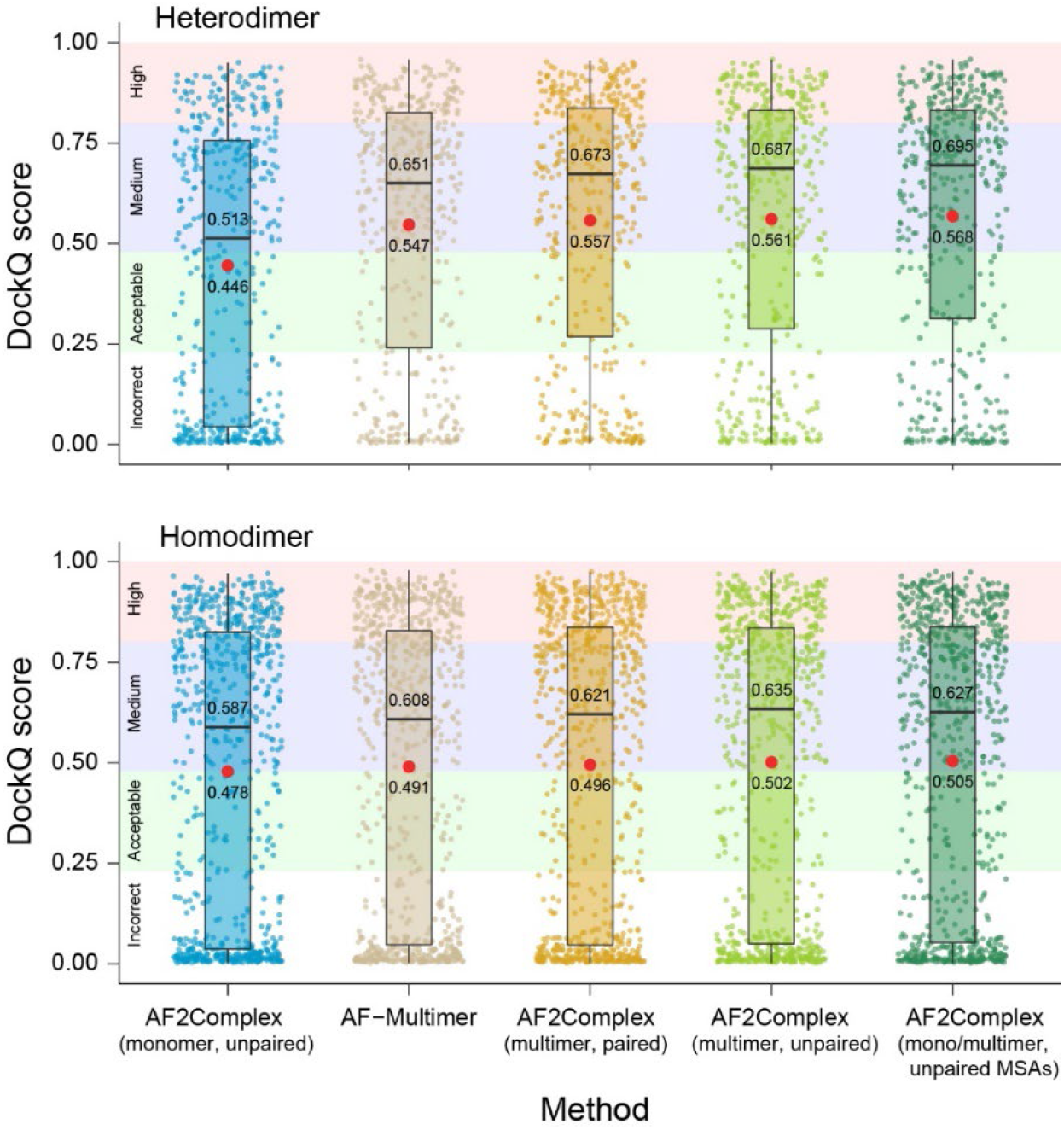
Comparison of AF-Multimer and various AF2Complex strategies on the Dimer1193 set. Evaluation scores of the top 1 ranked models for 440 heterodimers (top panel) and 753 homodimers (bottom) are scattered small circles and color-coded for different methods. Types of AF deep learning models used are denoted as “monomer” or “multimer”. MSA styles are indicated by “paired” or “unpaired”. The AF2Complex runs with paired MSAs used up to 20 recycles instead of the default 3 cycles in the AF-Multimer runs. Other runs also were carried out with up to 20 recycles. Black boxes and bars represent the second and third quartiles (25% to 75% ranked by the DockQ scores) and the medians of the distributions. Red stars represent the mean values. Numeric median and mean values are shown. The colored background panels indicate the regimes of high, medium, acceptable, and incorrect complex models.

On the 440 heterodimers, we see clear improvement in terms of DockQ score[5] evaluations with the AF-Multimer DL models. The mean/median of AF2Complex using monomer DL models on the heterodimers is 0.446/0.513, which was significantly improved to 0.547/0.651 by AF-Multimer in its default setting with three recycles and paired MSAs (*p*-value < 2.2 × 10^-16^, Wilcoxon signed-rank test, paired, one-tailed, n = 440, same test employed below). If we allow additional recycles up to 20 and use the same paired MSAs as in AF-Multimer, the numbers increase further to 0.557/0.673 ((*p*-value = 0.11). By using unpaired MSAs, the mean/median elevate again to 0.561/0.687 (*p*-value = 5×10^-3^). Finally, if we evaluate the top 1 ranked (according to the interface-score) models obtained with either monomer or multimer DL models and unpaired MSAs, we obtain the best performance at 0.568/0.695 (*p*-value = 6×10^-4^). In this latest strategy, about 77% of heterodimeric targets have their overall top ranked model at acceptable or better quality, 68% at medium or better quality, and 31% at high quality. Correspondingly, the mean interface RMSDs of the assessed models according to DockQ evaluation are 0.79, 1.37, and 1.95 Å. By using unpaired MSAs, AF2Complex yields statistically significantly better complex models than AF-Multimer on the heterodimers.

However, the improvement with the multimer DL models is relatively small with the 753 homodimers. Using the monomer DL models, AF2Complex yields a mean/median of 0.479/0.589, versus 0.491/0.608 by AF-multimer, and 0.504/0.627 by AF2Complex using monomer/multimer models and unpaired MSAs. The difference is statistically significant from AF2Complex monomer models to AF-Multimer (*p*-value = 9.8×10^-7^, Wilcoxon test, n = 753), but further improvement by AF2Complex is statistically insignificant according to the same Wilcoxon test.

We also note that the predictions on the homodimers are somewhat worse than those of the heterodimers. On average, the mean DockQ score is about 0.05 lower between the two dimer sets with the same method. Two observations may explain the difference.

First, although most homodimers exhibit the two-fold cyclic symmetry (C2), 56 (7%) of them are asymmetric and they invariable have a low model quality because the DL models usually yield symmetric models on a homodimer target. If we remove these asymmetric dimer targets, the mean of DockQ increase about 0.025.

Second, a homodimer may have alternative interaction poses that come from higher order symmetry, e.g., dihedral symmetry D2 that have two distinct interfaces. We strived to remove such cases in the benchmark set by considering only the dimers that have a global symmetry of homodimer according to the PDB annotation, that is, they are not part of a higher order symmetry in the considered PDB records. However, some proteins may still form higher order complexes that are in separate PDB records or even absent in the PDB. If we remove 46 (6%) cases with DockQ score < 0.23 and high Interface-score > 0.6 (only 8 (2%) such cases found in the heterodimer counterpart set), we obtain another 0.03 increase in the mean of the DockQ scores. These cases could be alternative docking poses that are not shown in the targeted experimental structures.

For instance, the target 5XBT contains a homodimer in C2 symmetry, but the same protein also appears as a homotetramer in a separate PDB record (5XBW), which exhibits D2 symmetry[6]. AF2Complex correctly predicted one interaction pose that appears in the D2 form but not the one in the C2 form. When we modeled the homotetramer form of this target with the multimer DL models and unpaired MSAs, we obtained a high-quality tetramer model with a TM-score of 0.92 compared to the experimental structure.

### AF-Multimer may yield unphysical structural models for large oligomers

Despite clear improvement made possible by the multimer DL models, they come with a limitation in that they could generate unphysical models with many atomic clashes at protein-protein interfaces. This effect is illustrated in Fig. S3a, where structural models of a homodimer target from Dimer1193 set are shown. This target has a long, disordered central segment (~200 residues) missing in the crystal structure, where the N- and C-terminal segments are found to fold into single ribokinase domains and two of them form a homodimer[7]. With the full sequence as the input, AF-Multimer generates a model (top 1 ranked), whereby the central segments overlap with each other and the ribokinase domains. The clashes are dramatically reduced in the top model by AF2Complex, using the same multimer DL models but unpaired MSAs. The most physical computational model comes from monomer DL models and unpaired MSAs. In this model, the central segments largely swing far away from the folded ribokinase domains. All three computational models displayed have a very high DockQ-score of ~0.95, and the ribokinase domains superimposed near perfectly with the experimental structures (Fig. S3b). This result occurs because the clashed regions are missing in the experimental structures, and therefore, ignored in the model evaluation. However, the model obtained with AF-Multimer default settings is unphysical and misleading.

To address this omission in model evaluation, we introduce a simple metric called the *interface clash indicator* χ, which is defined as the number of interface residues divided by the number of interface residue-residue contacts (see Methods). In the example above, the experimental structure has χ = 0.83, which is very close to χ = 0.81 for the monomer model, and 0.78 for the multimer model obtained with unpaired MSAs. By contrast, the AF-Multimer model has a χ value of 0.28, well below 0.5, which is the observed lower boundary of experimental structures.

The clashed interfaces are more often observed in large oligomers than dimers with multimer DL models. Fig. S3b shows the statistics of predicted models and the experimental structures of corresponding targets from the Oligomer562 set, each target with 3 or more monomers. None of experimental structures in this set had a χ value less than 0.6. Extensively intertwined structures, such as those that occur through domain-are like overwhelmed by clashing interface residues at χ < 0.6. The issue is somewhat alleviated if unpaired MSAs are used with the multimer models, reducing the percentage to 14%. The monomer DL models with unpaired input suffer this clash issue to a much-reduced degree, with about 1.3% of models at χ < 0.6. Note that these clashes cannot be eliminated by the AF2 relaxation procedure, which is a molecular dynamics minimization step barely moving protein backbone. Such large backbone movement is needed to remove the clashes. However, the relaxation could remove some minor side chain clashes observed within structures with higher χ values.

**Fig. S3.**
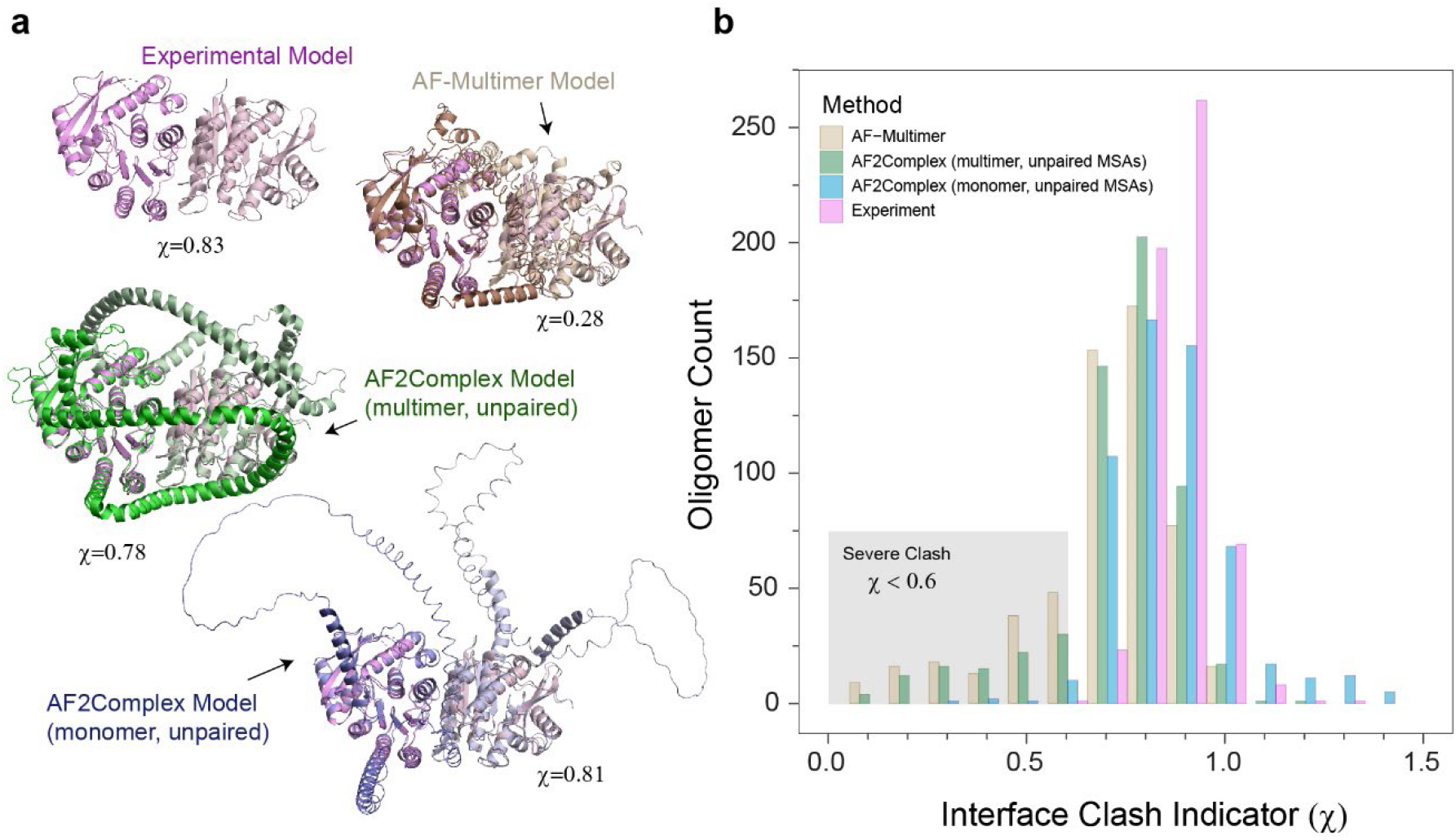
Analysis of residue clashes at protein-protein interface of predicted protein complex models. (**a**) An illustrative example from the Dimer1193 set is a pyridoxin/pyridoxal kinase (PdxK) from *Plasmodium falciparum*. An experimental structure (PDB code: 6SU9) revealed that the terminal domains of this target protein form a homodimer, but the central part of ~200 residues are missing in the crystal structure. The full dimer sequences were modeled by different methods and the top 1 ranked models are superimposed onto the experimental structure. The structural models are shown in the cartoon representation. (**b**) Histogram of the interface clash indicator on the Oligomer562 set. For each method, the overall top ranked models were analyzed. The regime where predicted structural models contains severe clashes are marked by a grey rectangle. Note that the AF2 relaxation step was not applied to these models. The relaxation can remove minor clashes caused by small side chain reorientations, but it cannot remove the clashes requiring significant backbone movements such as the example shown in (a).

Overall, the multimer DL models can generate higher quality oligomer models than monomer DL models. However, they may also yield unphysical models with many clashes at the interface, whereas the monomer DL models are much more physical. The issue may be alleviated by using unpaired MSAs, but not eliminated. We fully expect that DeepMind will address this issue with improved neural network models for multimeric complex prediction in the near future.

## Notes

https://github.com/FreshAirTonight/af2complex

https://zenodo.org/record/6084186

### Competing Interest Statement

The authors have declared no competing interest.

### Summary of Updates

1. Added support to AlphaFold-Multimer neural network models. 2. Performed an additional large-scale benchmark test. 3. Conducted additional modeling of the E. coli Ccm I system.

## References

1. Jumper, J., R. Evans, A. Pritzel, T. Green, M. Figurnov, O. Ronneberger, et al., Highly accurate protein structure prediction with AlphaFold. Nature, 2021. 596(7873): p. 583–589.

2. Tunyasuvunakool, K., J. Adler, Z. Wu, T. Green, M. Zielinski, A. žídek, et al., Highly accurate protein structure prediction for the human proteome. Nature, 2021. 596(7873): p. 590–596.

3. Skolnick, J., M. Gao, H. Zhou, and S. Singh, AlphaFold 2: Why It Works and Its Implications for Understanding the Relationships of Protein Sequence, Structure, and Function. Journal of Chemical Information and Modeling, 2021. 61(10): p. 4827–4831.

4. Marcotte, E.M., M. Pellegrini, H.-L. Ng, D.W. Rice, T.O. Yeates, and D. Eisenberg, Detecting Protein Function and Protein-Protein Interactions from Genome Sequences. Science, 1999. 285(5428): p. 751–753.

5. Keskin, Z., A. Gursoy, B. Ma, and R. Nussinov, Principles of protein-protein interactions: What are the preferred ways for proteins to interact? Chem. Rev., 2008. 108(4): p. 1225–1244.

6. Gao, M. and J. Skolnick, Structural space of protein-protein interfaces is degenerate, close to complete, and highly connected. Proc. Natl. Acad. Sci. U. S. A., 2010. 107(52): p. 22517–22522.

7. Baek, M., F. DiMaio, I. Anishchenko, J. Dauparas, S. Ovchinnikov, G.R. Lee, et al., Accurate prediction of protein structures and interactions using a three-track neural network. Science, 2021. 373(6557): p. 871–876.

8. Vakser, I.A., Protein-protein docking: from interaction to interactome. Biophys J, 2014. 107(8): p. 1785–1793.

9. Kozakov, D., R. Brenke, S.R. Comeau, and S. Vajda, PIPER: An FFT-based protein docking program with pairwise potentials. Proteins, 2006. 65(2): p. 392–406.

10. Dominguez, C., R. Boelens, and A. Bonvin, HADDOCK: A protein-protein docking approach based on biochemical or biophysical information. J. Am. Chem. Soc., 2003. 125(7): p. 1731–1737.

11. Chen, R., L. Li, and Z.P. Weng, ZDOCK: An initial-stage protein-docking algorithm. Proteins-Structure Function and Genetics, 2003. 52(1): p. 80–87.

12. Aloy, P., B. Bottcher, H. Ceulemans, C. Leutwein, C. Mellwig, S. Fischer, et al., Structure-based assembly of protein complexes in yeast. Science, 2004. 303(5666): p. 2026–2029.

13. Chen, H.L. and J. Skolnick, M-TASSER: An algorithm for protein quaternary structure prediction. Biophys. J., 2008. 94(3): p. 918–928.

14. Zhang, Q.C., D. Petrey, R. Norel, and B.H. Honig, Protein interface conservation across structure space. Proc. Natl. Acad. Sci. U. S. A., 2010. 107(24): p. 10896–10901.

15. Keskin, O., R. Nussinov, and A. Gursoy, PRISM: protein-protein interaction prediction by structural matching. Methods Mol Biol, 2008. 484: p. 505–21.

16. Mukherjee, S. and Y. Zhang, Protein-Protein Complex Structure Predictions by Multimeric Threading and Template Recombination. Structure, 2011. 19(7): p. 955–966.

17. Szurmant, H. and M. Weigt, Inter-residue, inter-protein and inter-family coevolution: bridging the scales. Curr Opin Struct Biol, 2018. 50: p. 26–32.

18. Zhou, T.-m., S. Wang, and J. Xu, Deep learning reveals many more inter-protein residue-residue contacts than direct coupling analysis. bioRxiv, 2017: p. 240754.

19. Ko, J. and J. Lee, Can AlphaFold2 predict protein-peptide complex structures accurately? bioRxiv, 2021: p. 2021.07.27.453972.

20. Mirdita, M., S. Ovchinnikov, and M. Steinegger, ColabFold - Making protein folding accessible to all. bioRxiv, 2021: p. 2021.08.15.456425.

21. Bryant, P., G. Pozzati, and A. Elofsson, Improved prediction of protein-protein interactions using AlphaFold2. bioRxiv, 2021: p. 2021.09.15.460468.

22. Ghani, U., I. Desta, A. Jindal, O. Khan, G. Jones, S. Kotelnikov, et al., Improved Docking of Protein Models by a Combination of Alphafold2 and ClusPro. bioRxiv, 2021: p. 2021.09.07.459290.

23. Uetz, P., L. Giot, G. Cagney, T.A. Mansfield, R.S. Judson, J.R. Knight, et al., A comprehensive analysis of protein-protein interactions in Saccharomyces cerevisiae. Nature, 2000. 403(6770): p. 623–627.

24. Butland, G., J.M. Peregrín-Alvarez, J. Li, W. Yang, X. Yang, V. Canadien, et al., Interaction network containing conserved and essential protein complexes in Escherichia coli. Nature, 2005. 433(7025): p. 531–537.

25. Arifuzzaman, M., M. Maeda, A. Itoh, K. Nishikata, C. Takita, R. Saito, et al., Large-scale identification of protein–protein interaction of Escherichia coli K-12. Genome Research, 2006. 16(5): p. 686–691.

26. Yu, H., P. Braun, M.A. Yildirim, I. Lemmens, K. Venkatesan, J. Sahalie, et al., High-quality binary protein interaction map of the yeast interactome network. Science, 2008. 322(5898): p. 104–110.

27. Gong, W., A. Guerler, C. Zhang, E. Warner, C. Li, and Y. Zhang, Integrating Multimeric Threading With High-throughput Experiments for Structural Interactome of Escherichia coli. Journal of Molecular Biology, 2021. 433(10): p. 166944.

28. Cong, Q., I. Anishchenko, S. Ovchinnikov, and D. Baker, Protein interaction networks revealed by proteome coevolution. Science, 2019. 365(6449): p. 185–189.

29. Humphreys, I.R., J. Pei, M. Baek, A. Krishnakumar, I. Anishchenko, S. Ovchinnikov, et al., Computed structures of core eukaryotic protein complexes. Science, 2021. 374(6573): p. eabm4805.

30. Koonin, E.V., Orthologs, paralogs, and evolutionary genomics. Annu Rev Genet, 2005. 39: p. 309–38.

31. Gao, M. and J. Skolnick, iAlign: a method for the structural comparison of protein-protein interfaces. Bioinformatics, 2010. 26(18): p. 2259–2265.

32. Hu, P., S.C. Janga, M. Babu, J.J. Díaz-Mejía, G. Butland, W. Yang, et al., Global Functional Atlas of Escherichia coli Encompassing Previously Uncharacterized Proteins. PLOS Biology, 2009. 7(4): p. e1000096.

33. Sanders, C., S. Turkarslan, D.W. Lee, and F. Daldal, Cytochrome c biogenesis: the Ccm system. Trends Microbiol, 2010. 18(6): p. 266–74.

34. Kranz, R.G., C. Richard-Fogal, J.S. Taylor, and E.R. Frawley, Cytochrome c biogenesis: mechanisms for covalent modifications and trafficking of heme and for heme-iron redox control. Microbiol Mol Biol Rev, 2009. 73(3): p. 510–28, Table of Contents.

35. Evans, R., M. O’Neill, A. Pritzel, N. Antropova, A. Senior, T. Green, et al., Protein complex prediction with AlphaFold-Multimer. bioRxiv, 2021: p. 2021.10.04.463034.

36. Ozden, B., A. Kryshtafovych, and E. Karaca, Assessment of the CASP14 assembly predictions. Proteins, 2021.

37. Lensink, M.F., G. Brysbaert, T. Mauri, N. Nadzirin, S. Velankar, R.A.G. Chaleil, et al., Prediction of protein assemblies, the next frontier: The CASP14-CAPRI experiment. Proteins, 2021.

38. Humphrey, W., A. Dalke, and K. Schulten, VMD: visual molecular dynamics. Journal of Molecular Graphics, 1996. 14(1): p. 33–38.

39. Dunce, J.M., L.J. Salmon, and O.R. Davies, Structural basis of meiotic chromosome synaptic elongation through hierarchical fibrous assembly of SYCE2-TEX12. Nature Structural & Molecular Biology, 2021. 28(8): p. 681–693.

40. Zhang, Y. and J. Skolnick, Scoring function for automated assessment of protein structure template quality. Proteins, 2004. 57(4): p. 702–710.

41. Gao, M. and J. Skolnick, New benchmark metrics for protein-protein docking methods. Proteins, 2011. 79(5): p. 1623–1634.

42. Kozakov, D., D.R. Hall, B. Xia, K.A. Porter, D. Padhorny, C. Yueh, et al., The ClusPro web server for protein–protein docking. Nature Protocols, 2017. 12(2): p. 255–278.

43. Basu, S. and B. Wallner, DockQ: A Quality Measure for Protein-Protein Docking Models. PLOS ONE, 2016. 11(8): p. e0161879.

44. Feissner, R.E., C.L. Richard-Fogal, E.R. Frawley, and R.G. Kranz, ABC transporter-mediated release of a haem chaperone allows cytochrome c biogenesis. Mol Microbiol, 2006. 61(1): p. 219–31.

45. San Francisco, B. and R.G. Kranz, Interaction of HoloCcmE with CcmF in Heme Trafficking and Cytochrome c Biosynthesis. Journal of Molecular Biology, 2014. 426(3): p. 570–585.

46. San Francisco, B., M.C. Sutherland, and R.G. Kranz, The CcmFH complex is the system I holocytochrome c synthetase: engineering cytochrome c maturation independent of CcmABCDE. Molecular Microbiology, 2014. 91(5): p. 996–1008.

47. Verissimo, A.F., B. Khalfaoui-Hassani, J. Hwang, S. Steimle, N. Selamoglu, C. Sanders, et al., The thioreduction component CcmG confers efficiency and the heme ligation component CcmH ensures stereo-specificity during cytochrome c maturation. J Biol Chem, 2017. 292(32): p. 13154–13167.

48. Sutherland, M.C., J.M. Jarodsky, S. Ovchinnikov, D. Baker, and R.G. Kranz, Structurally Mapping Endogenous Heme in the CcmCDE Membrane Complex for Cytochrome c Biogenesis. J Mol Biol, 2018. 430(8): p. 1065–1080.

49. Richard-Fogal, C. and R.G. Kranz, The CcmC:heme:CcmE complex in heme trafficking and cytochrome c biosynthesis. Journal of molecular biology, 2010. 401(3): p. 350–362.

50. Brausemann, A., L. Zhang, L. Ilcu, and O. Einsle, Architecture of the membrane-bound cytochrome c heme lyase CcmF. Nat Chem Biol, 2021. 17(7): p. 800–805.

51. Mendez, D.L., E.P. Lowder, D.E. Tillman, M.C. Sutherland, A.L. Collier, M.J. Rau, et al., Cryo-EM of CcsBA reveals the basis for cytochrome c biogenesis and heme transport. Nature Chemical Biology, 2022. 18(1): p. 101–108.

52. Xu, J., M. McPartlon, and J. Li, Improved protein structure prediction by deep learning irrespective of co-evolution information. Nature Machine Intelligence, 2021. 3(7): p. 601–609.

53. The UniProt, C., UniProt: a worldwide hub of protein knowledge. Nucleic Acids Research, 2019. 47(D1): p. D506–D515.

54. Mitchell, A.L., A. Almeida, M. Beracochea, M. Boland, J. Burgin, G. Cochrane, et al., MGnify: the microbiome analysis resource in 2020. Nucleic Acids Research, 2020. 48(D1): p. D570–D578.

55. Mirdita, M., L. von den Driesch, C. Galiez, M.J. Martin, J. Söding, and M. Steinegger, Uniclust databases of clustered and deeply annotated protein sequences and alignments. Nucleic acids research, 2017. 45(D1): p. D170–D176.

56. Steinegger, M., M. Mirdita, and J. Söding, Protein-level assembly increases protein sequence recovery from metagenomic samples manyfold. Nature Methods, 2019. 16(7): p. 603–606.

57. Berman, H.M., J. Westbrook, Z. Feng, G. Gilliland, T.N. Bhat, H. Weissig, et al., The Protein Data Bank. Nucleic Acids Research, 2000. 28(1): p. 235–242.

58. Mukherjee, S. and Y. Zhang, MM-align: a quick algorithm for aligning multiple-chain protein complex structures using iterative dynamic programming. Nucleic Acids Research, 2009. 37(11): p. e83.

59. Wu, C.H., R. Apweiler, A. Bairoch, D.A. Natale, W.C. Barker, B. Boeckmann, et al., The Universal Protein Resource (UniProt): an expanding universe of protein information. Nucleic Acids Research, 2006. 34: p. D187–D191.

60. Clarke, T.A., G.L. Kemp, J.H.V. Wonderen, R.-M.A.S. Doyle, J.A. Cole, N. Tovell, et al., Role of a Conserved Glutamine Residue in Tuning the Catalytic Activity of Escherichia coli Cytochrome c Nitrite Reductase. Biochemistry, 2008. 47(12): p. 3789–3799.

61. Nivón, L.G., R. Moretti, and D. Baker, A Pareto-Optimal Refinement Method for Protein Design Scaffolds. PLOS ONE, 2013. 8(4): p. e59004.

62. Sing, T., O. Sander, N. Beerenwinkel, and T. Lengauer, ROCR: visualizing classifier performance in R. Bioinformatics, 2005. 21(20): p. 3940–3941.

63. Gao, M., P. Lund-Andersen, A. Morehead, S. Mahmud, C. Chen, X. Chen, et al. High-Performance Deep Learning Toolbox for Genome-Scale Prediction of Protein Structure and Function. in 2021 IEEE/ACM Workshop on Machine Learning in High Performance Computing Environments (MLHPC). 2021.

64. Gao, M., M. Coletti, R.B. Davidson, R. Prout, S. Abraham, B. Hernandez, et al., Proteome-scale Deployment of Protein Structure Prediction Workflows on the Summit Supercomputer. arXiv preprint arXiv:2201.10024, 2022.

## Supplementary References

1. Evans, R., M. O’Neill, A. Pritzel, N. Antropova, A. Senior, T. Green, et al., Protein complex prediction with AlphaFold-Multimer. bioRxiv, 2021: p. 2021.10.04.463034.

2. Ozden, B., A. Kryshtafovych, and E. Karaca, Assessment of the CASP14 assembly predictions. Proteins, 2021.

3. Jumper, J., R. Evans, A. Pritzel, T. Green, M. Figurnov, O. Ronneberger, et al., Applying and improving AlphaFold at CASP14. Proteins: Structure, Function, and Bioinformatics, 2021. 89(12): p. 1711–1721.

4. Gao, M. and J. Skolnick, iAlign: a method for the structural comparison of protein-protein interfaces. Bioinformatics, 2010. 26(18): p. 2259–2265.

5. Basu, S. and B. Wallner, DockQ: A Quality Measure for Protein-Protein Docking Models. PLOS ONE, 2016. 11(8): p. e0161879.

6. Wang, F., Q. He, J. Yin, S. Xu, W. Hu, and L. Gu, BrlR from Pseudomonas aeruginosa is a receptor for both cyclic di-GMP and pyocyanin. Nature Communications, 2018. 9(1): p. 2563.

7. Gao, K., W. Wang, T. Kronenberger, C. Wrenger, and M.R. Groves, The Crystal Structure of the Plasmodium falciparum PdxK Provides an Experimental Model for Pro-Drug Activation. Crystals, 2019. 9(10): p. 534.

